# RNP-MaP: In-cell analysis of protein interaction networks defines functional hubs in RNA

**DOI:** 10.1101/2020.02.07.939108

**Authors:** Chase A. Weidmann, Anthony M. Mustoe, Parth B. Jariwala, J. Mauro Calabrese, Kevin M. Weeks

**Affiliations:** Department of Chemistry, University of North Carolina, Chapel Hill NC 27599-3290; Lineberger Comprehensive Cancer Center, University of North Carolina, Chapel Hill, NC 27599; Department of Pharmacology, University of North Carolina, Chapel Hill, NC 27599, USA

## Abstract

RNAs interact with networks of proteins to form complexes (RNPs) that govern many biological processes, but these networks are currently impossible to examine in a comprehensive way. We developed a live-cell chemical probing strategy for mapping protein interaction networks in any RNA with single-nucleotide resolution. This RNP-MaP strategy (<u>RNP</u> network analysis by <u>m</u>ut<u>a</u>tional <u>p</u>rofiling) simultaneously detects binding by and cooperative interactions involving multiple proteins with single RNA molecules. RNP-MaP revealed that two structurally related, but sequence-divergent noncoding RNAs, RNase P and RMRP, share nearly identical RNP networks and, further, that protein interaction network hubs identify function-critical sites in these RNAs. RNP-MaP identified numerous protein interaction networks within the XIST long noncoding RNA that are conserved between mouse and human RNAs and distinguished communities of proteins that network together on XIST. RNP-MaP data show that the Xist E region is densely networked by protein interactions and that PTBP1, MATR3, and TIA1 proteins each interface with the XIST E region via two distinct interaction modes; and we find that the XIST E region is sufficient to mediate RNA foci formation in cells. RNP-MaP will enable discovery and mechanistic analysis of protein interaction networks across any RNA in cells.

## INTRODUCTION

RNA-protein complexes (RNPs) govern both mRNA regulation and noncoding RNA (ncRNA) function^1, 2^. Understanding how RNPs assemble and function, involving both one RNA–one protein interactions and multi-component interaction networks, is thus critical for characterizing many biological processes and mechanisms. To date, biochemical approaches have defined in detail protein interactions required for a number of RNP assemblies^3^, and high-resolution structural approaches^4–6^ have revolutionized our understanding of small and large RNP architectures. Nonetheless, it remains challenging to characterize RNP assemblies and their interacting networks within living systems.

Current methods for characterizing RNPs in live cells suffer from several limitations. Crosslinking and immunoprecipitation (CLIP) approaches have enabled identification of binding sites for individual RNA-binding proteins transcriptome-wide^7^. However, CLIP suffers from strong experimental biases^8^, is limited in binding-site resolution, assesses only a single protein at a time, and requires a unique antibody or exogenous tag. Methods that focus on cataloging RNA-binding proteins, like mass spectrometry, cannot simultaneously locate protein-binding sites on RNA nor easily prioritize proteins in terms of function^9^. These limitations obscure the importance of binding by multiple proteins to individual RNAs. Major unaddressed challenges are: (1) How do multiple RNA-binding proteins cooperate on an RNA to form networks, and (2) Which protein interaction networks are the most critical for function of an individual RNA?

Here we describe an experimentally concise strategy that locates sites of protein interaction on any targeted RNA in live cells with nucleotide resolution, reports on multi-protein interaction networks present on single RNA molecules, and identifies network hubs most integral to the function of an RNP. We validated this approach, termed ribo<u>n</u>ucleo<u>p</u>rotein network analysis by <u>m</u>ut<u>a</u>tional <u>p</u>rofiling or RNP-MaP, on small and large RNPs with known architectures. RNP-MaP was then used to define critical functional regions of the XIST long noncoding RNA (lncRNA), to categorize XIST-binding proteins into distinct interactive groups, and to identify features of the functionally important XIST E region. RNP-MaP will be widely useful for characterizing RNP biology, particularly in defining functionally critical and disease-relevant domains in large messenger and non-coding RNAs.

## RESULTS

### RNP-MaP Validation

To comprehensively map protein interaction networks of an RNA of interest, we identified a cell-permeable reagent, NHS-diazirine (SDA), that can rapidly label RNA nucleotides at sites of protein binding. SDA has two reactive moieties: a succinimidyl ester and a diazirine (Fig. 1a). Succinimidyl esters react to form amide bonds with amines such as those found in lysine side chains. When photoactivated with long-wavelength ultraviolet light (UV-A, 365 nm), diazirines form carbene or diazo intermediates^10^, which are broadly reactive to nucleotide sugar and base moieties. The two-step reaction of SDA thus crosslinks protein lysine residues with RNA with a distance governed by SDA linker length (4 Å) and lysine flexibility (6 Å). Lysine is one of the most prevalent amino acids in RNA-binding domains^11^, and photo-intermediate lifetimes are short; thus, SDA is expected to crosslink short-range RNA-protein interactions relatively independently of local RNA structure or protein properties. To perform crosslinking, live cells are treated with SDA for a short time and then exposed to UV-A light.

**Figure 1.**
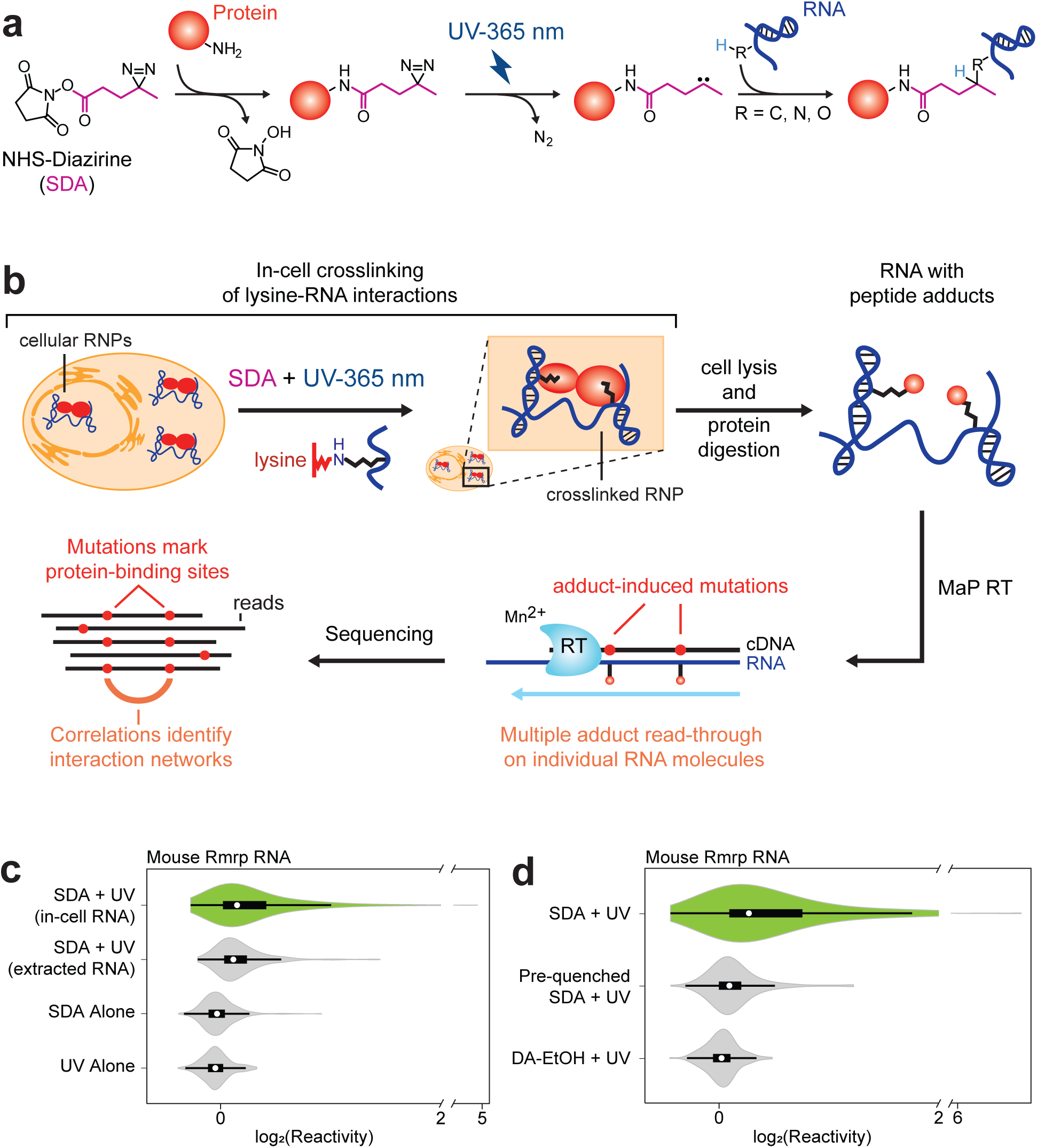
RNP-MaP strategy for probing RNA-protein interaction networks in cells. (a) Scheme for selective chemical crosslinking of protein to RNA by SDA. (b) Workflow of the RNP-MaP experiment. (c) Relative mutation rate over background [log_2_(reactivity)] of Rrmp RNA with and without SDA, UV, and cellular proteins. (d) log_2_(reactivity) of Rmrp RNA in the presence of SDA, pre-quenched SDA, and DA-EtOH. In violin plots shown in panels C and D, circles indicate medians, box limits indicate the first and third quartiles, whiskers extend 1.5 times the interquartile range, and smoothed polygons show data density estimates and extend to extreme values.

To detect SDA-mediated RNA-protein crosslinks, we use the MaP reverse transcription technology (Fig. 1b). SDA-treated cells are lysed and crosslinked proteins are digested to short peptide adducts. Adduct-containing RNA is then used as a template for relaxed-fidelity reverse transcription^12, 13^. Under MaP conditions, reverse transcriptase reads through adduct-containing nucleotides and incorporates non-templated nucleotides into the product DNA at the site of RNA-protein crosslinks. Importantly, because transcription reads through adducts, RNP-MaP detects multiple protein crosslinks that co-occur on single RNA molecules (Fig. 1b). Sequencing the DNA product and locating sites of mutation thus reveals two key features of an RNA-protein complex: RNP-MaP adducts at individual nucleotides report locations of protein binding, and correlated crosslinking across multiple nucleotides reveals higher-order protein interaction networks.

In experiments with U1 small nuclear RNA (snRNA), ribonuclease P RNA (RNase P), and the RNA component of mitochondrial RNA processing endoribonuclease (Rmrp) in mouse embryonic stem cells, the RNP-MaP signal was strictly dependent on treatment of cells with both UV and SDA (Fig. 1c **and** Fig. S1). To confirm that RNP-MaP adducts reflect bound proteins, we performed RNP-MaP on RNA extracted from cells (removing protein), which yielded substantially reduced adduct formation rates (Fig. 1c **and** Fig. S1). Use of either a diazirine ethanol compound (DA-EtOH), which has no lysine-reactive group, or SDA with a pre-quenched NHS-ester eliminated adduct formation, indicating that diazirine reactivity with RNA requires conjugation with lysine residues (Fig. 1d). In sum, RNP-MaP measures crosslinking events that are strictly dependent on SDA and UV dosage and that only occur in the presence of both RNA and cellular proteins.

We quantified distances between lysine residues and SDA-reactive sites of human U1, RNase P, and 18S and 28S ribosomal RNAs by comparison to high-resolution structures^4, 6, 14^. From these experiments, we derived nucleotide-specific normalization factors for calculating reactivity thresholds relative to experimental medians (Fig. S2). These normalization factors can be applied to RNP-MaP experiments on RNAs whose structure and protein partners are unknown, enabling de novo identification of protein-bound nucleotides with high confidence (Fig. S2). Nucleotides whose SDA-enhanced mutation rates (reactivities) exceed computed reactivity thresholds are called “RNP-MaP sites”. Reactivities were highly reproducible, nucleotides with higher reactivities were shorter distances from lysine amines, RNP-MaP sites were observed in both paired and unpaired regions of RNA and independently of nucleotide identity, and nucleotides distant from bound proteins rarely passed reactivity thresholds (Fig. S3). RNP-MaP sites identified in Rmrp and RNase P RNAs agreed with orthogonal ΔSHAPE chemical probing^15, 16^ and photo-lysine metabolic labeling^17^ (Fig. S4). These experiments demonstrated that RNP-MaP identifies short-distance crosslinks between an RNA and bound proteins.

**Figure 2.**
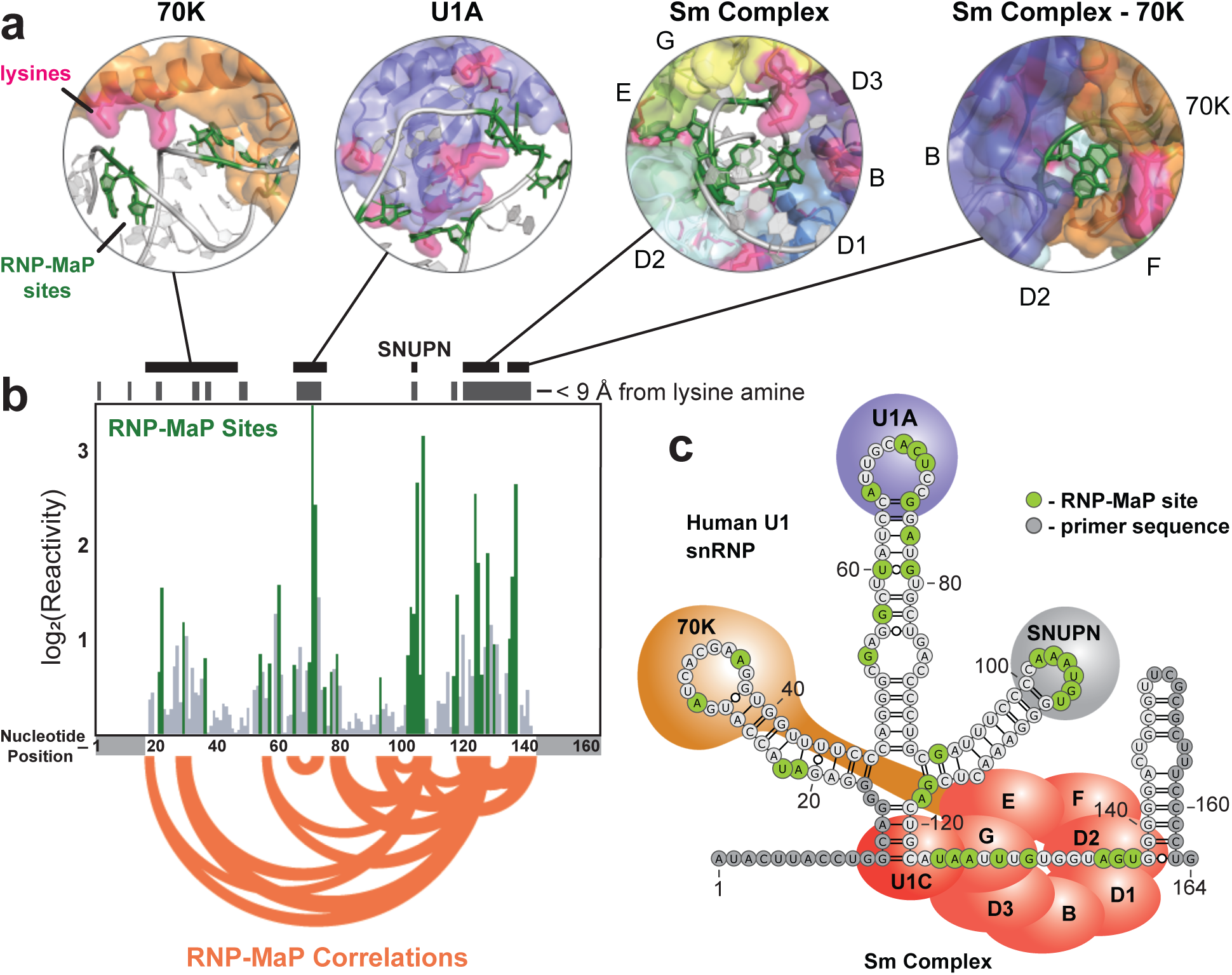
RNP-MaP correctly defines protein interaction networks in the U1 snRNP. (a) Structures surrounding lysine-RNA crosslinking sites (structures from 3CW1 and 4PKD^14^). (b) Bar graph of log_2_(average reactivity) from two replicate experiments on U1 snRNA in HEK293 cells. RNP-MaP sites passing thresholds are shown in green. Nucleotides within 9 Å of a lysine amine are highlighted with gray boxes above, and black boxes indicate exact locations of structures represented in panel A. RNP-MaP correlations (top 10% in mutual information strength) are shown as orange arcs. Nucleotides that overlap amplification primers (light gray boxes below) are not observable by RNP-MaP. (c) Secondary structure model of the human U1 RNP emphasizing relative protein positions, RNP-MaP sites, and primer regions.

**Figure 3.**
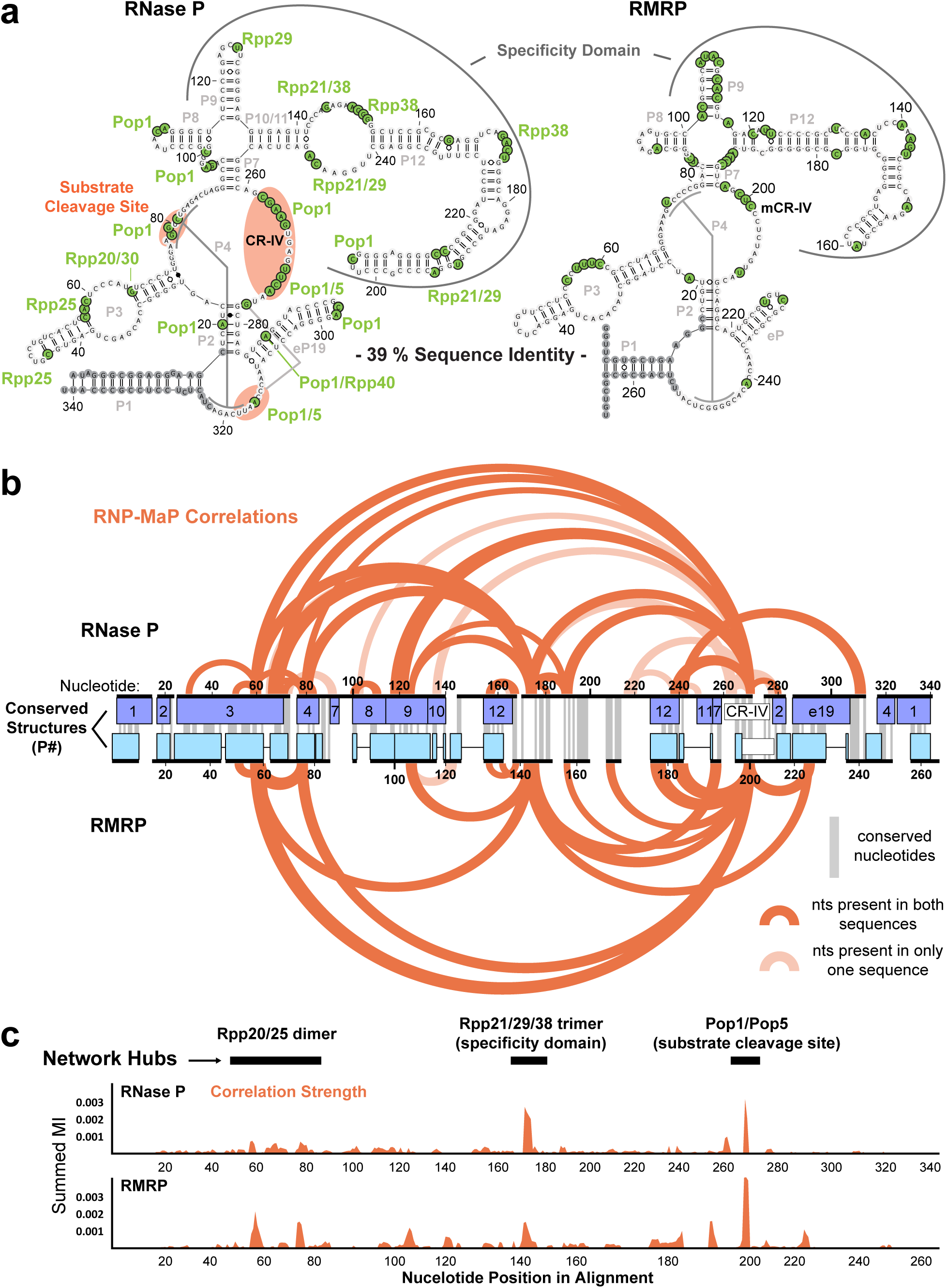
RNP-MaP reveals conserved protein interaction networks in RNase P and RMRP RNAs. (a) Secondary structure of human RNase P and RMRP RNAs with observed RNP-MaP sites identified in experiments in HEK293 cells highlighted in green. Proteins proximal to each site (judged by nearest lysine) are labeled. Important functional domains are indicated, and conserved base-paired structural regions (P#) are labeled in gray. (b) RNP-MaP correlations for human RNase P and RMRP, plotted on a structure-based sequence alignment. Conserved nucleotide positions and structural domains (P#) are labeled. Correlations shown are for the top 10% of mutual information strength. Correlations corresponding to linkages between nucleotides that are present only in one of the RNAs are shown in light shading. (c) The total strength of correlations (by mutual information, MI) at each nucleotide for the human RNase P and RMRP RNAs plotted on the same structure-based alignment. Protein interaction network hubs are labeled.

**Figure 4.**
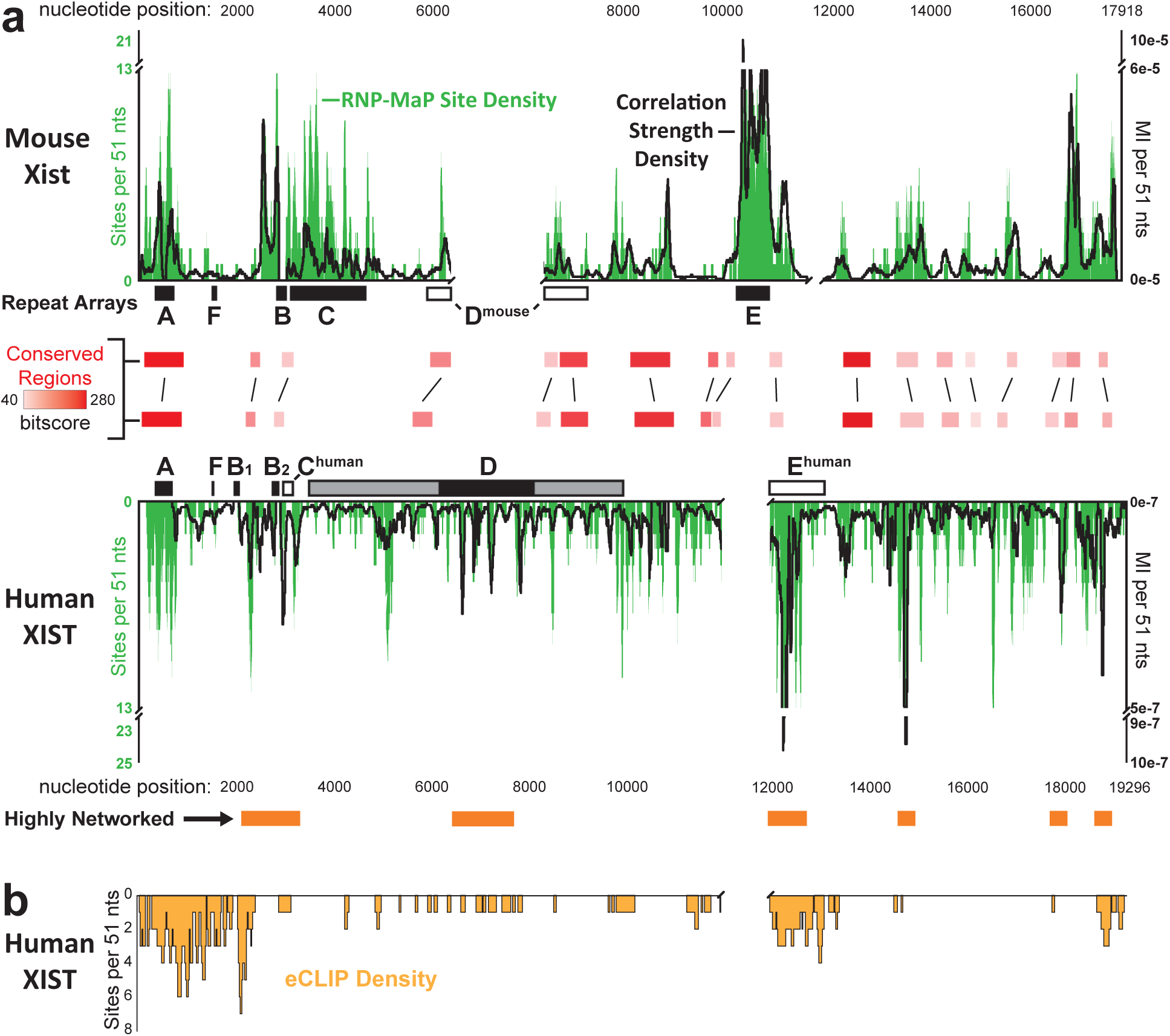
RNP-MaP identifies conserved protein interaction networks in the XIST lncRNA. (a) The density of RNP-MaP sites (green, the total number of RNP-MaP sites per 51-nt window, left axis) across the entirety of mouse Xist (top) and human XIST (bottom). Black lines indicate correlation strength densities (depth-normalized mutual information, MI, per 51-nt window, right axis). Highly networked regions within XIST are indicated with orange boxes below graph. Regions with conserved local sequence alignments are highlighted with red shading. Rectangles labeled A-F show locations of tandem repeat arrays within each RNA; filled rectangles denote well-defined repeat sequences, and open rectangles indicate regions identified as homologous in this work, as inferred by RNP-MaP similarity. The D region in human XIST contains a distinct repeat core (black) flanked by more degenerate repetitive elements (gray). Gaps were introduced to align RNAs by conservation and RNP-MaP similarity. Experiments were performed in SM33 and HEK293 cells. (b) Density of eCLIP sites along the human XIST RNA shown as the number of eCLIP sites within 51-nt windows. Based on 151 total sites from 30 proteins mapped reproducibly to XIST in K562 cells^37^.

### RNP-MaP defines protein interaction networks in the U1 snRNP

When RNP-MaP was used to examine protein interaction networks in the U1 snRNP in human HEK293 cells, we observed that RNP-MaP sites clustered in regions of U1 RNA known to bind proteins^14, 18^ (Fig. 2a, 2b). Notably, RNP-MaP sites occurred at all four ribonucleotides and in both single-stranded and base-paired regions (Fig. 2c), consistent with non-selective diazirine reactivity. The majority of RNP-MaP sites were within 9 Å of a lysine residue (Fig. 2b). We identified additional RNP-MaP sites in U1 stem loop 2 that, although close to the U1A binding site, do not correspond to known interaction sites (Fig. 2c). Only one of two RNA recognition motifs (RRMs) within the U1A protein is visualized in high-resolution structures^14^, and the RNP-MaP sites that extend outside of the known recognition site may reflect binding by the other U1A RRM or an unidentified protein component of the U1 snRNP.

Our RNP-MaP data also revealed multiple sites that were crosslinked in a correlated manner, reflecting RNA-mediated through-space protein-protein communication, in networks consistent with the known architecture of the U1 snRNP complex (Fig. 2b, 2c). Pairs of RNA sites exhibiting statistically significant co-mutations, termed RNP-MaP correlations, were identified using an established G-test framework^19^ (Fig. S2 **and** Fig. S5). The highest density and strength of correlations involved nucleotides bound by the Sm protein complex (Fig. 2b **and** Fig. S5), whose initial loading onto U1 is necessary for the maturation of the snRNP^20^. The second-highest density and strength of correlations were observed at the binding site for SNUPN protein, which recognizes Sm-bound U1 RNA and imports the RNP into the nucleus where it ultimately functions^21^. RNP-MaP correlations also reflect the long-distance network interactions between 70K and the Sm complex important for U1 snRNP assembly^22^. Only a few weak correlations were observed between the Sm core and U1A, consistent with the observation that U1A binds independently and that the stem-loop to which it binds is expendable for splicing activity^23^. Together, our data show that RNP-MaP correctly reveals protein interaction networks, pinpoints the central hubs of these networks, and identifies interactions important for assembly and function in the U1 snRNP.

**Figure 5.**
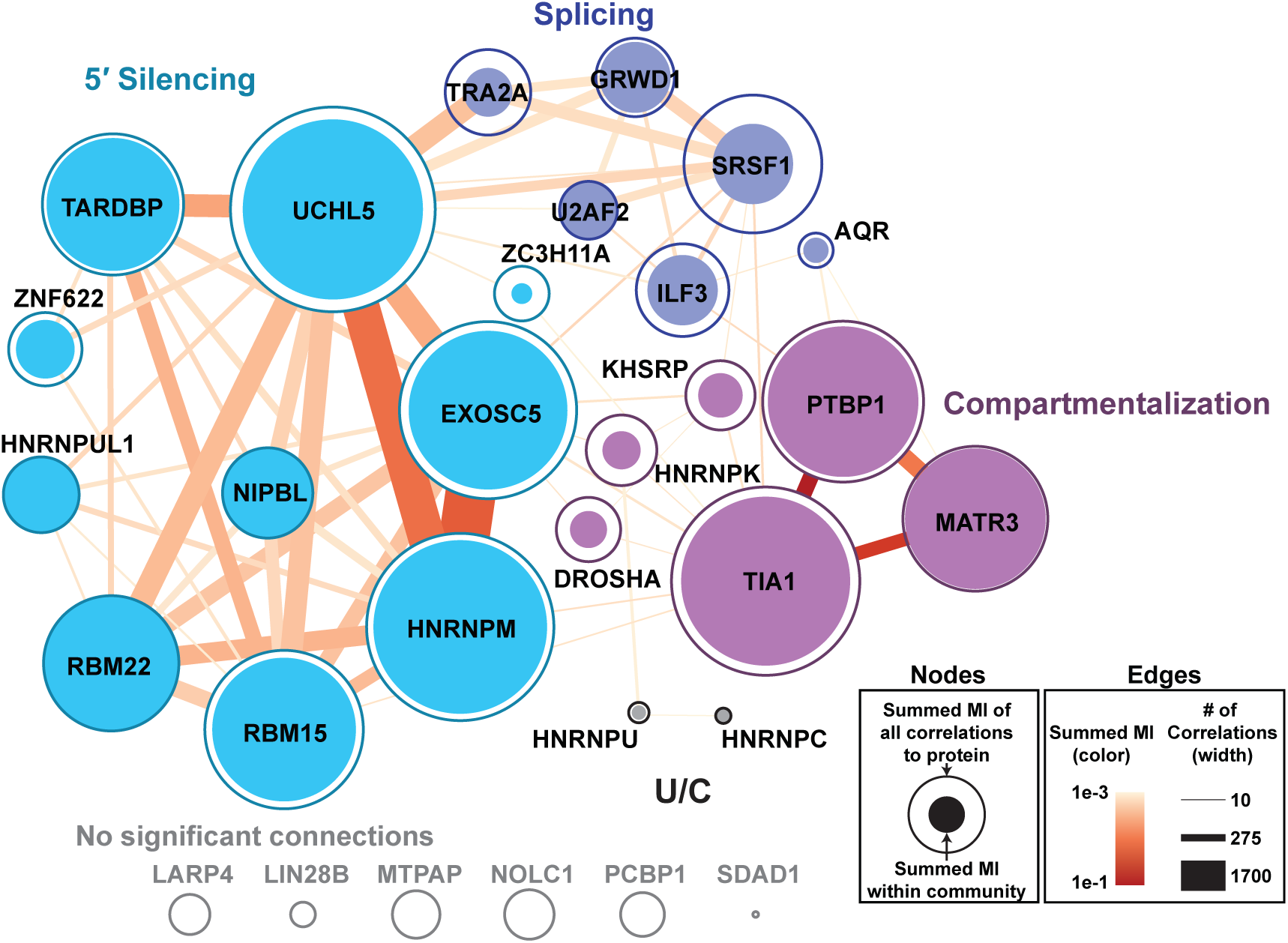
Communities of XIST-binding proteins. Network graph of protein-bound sites (nodes) from eCLIP data on human XIST RNA linked by RNP-MaP correlations (edges). Relative summed mutual information for correlations linking each protein node to other proteins in its community or to all proteins are shown as filled and open circles, respectively (see key). Summed mutual information of all correlations in an edge and total number of correlations in each edge are indicated by edge color and width, respectively.

### RNP-MaP reveals conserved protein interaction networks in RNase P and RMRP RNAs

RNase P and RMRP are structurally related, but sequence divergent, non-coding RNAs that bind overlapping sets of proteins to form RNP endonucleases that cleave distinct substrates^6, 19, 24^. RNP-MaP revealed that, despite differing sequences and substrates, the two RNPs have nearly identical protein interaction networks. We compared RNP-MaP sites to known protein interactions within RNase P^6^ and identified binding regions for 9 of the 10 RNase P interaction partners (Fig. 3a). Only Rpp14 binding was not detected, an absence explained by its limited interface with the RNA (of only eight amino acids, one of which is lysine). Comprehensive atomic resolution structural data does not exist for RMRP; however, the matching patterns of RNP-MaP sites for RNase P and RMRP suggest that most protein interaction sites outside of their specificity domains are shared. Relative locations of RNP-MaP sites are also conserved between human and mouse homologs of RNase P and RMRP (Fig. S6).

**Figure 6.**
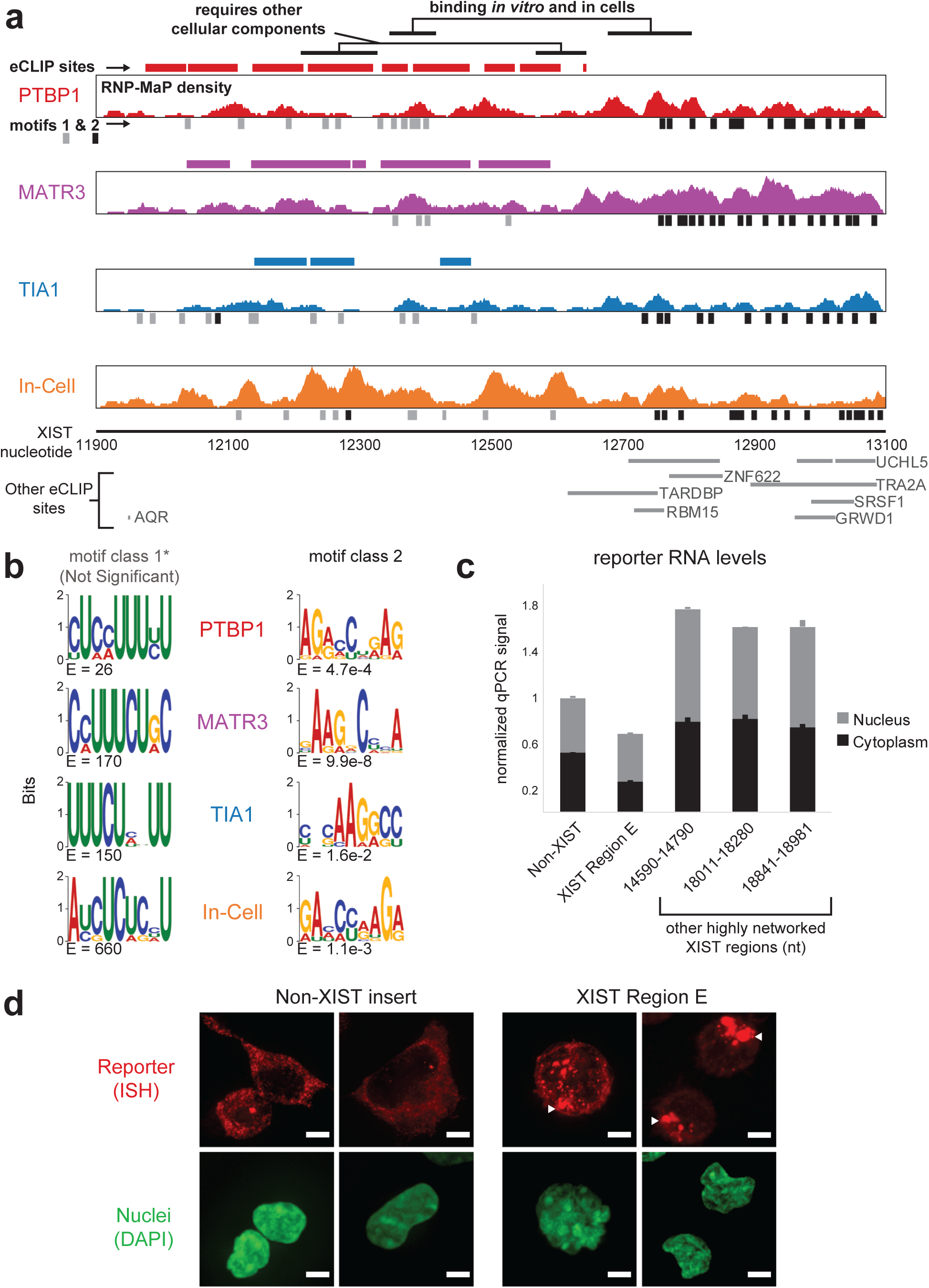
PTBP1, MATR3, and TIA1 recognize XIST E region via two binding modes. (a) Density of RNP-MaP sites on human XIST from experiments performed using an RNA spanning the E region and the indicated recombinant proteins (red, purple, blue). Experiment performed on native XIST in HEK293 cells is shown in orange. Locations of eCLIP sites and RNP-MaP enriched motifs corresponding to each protein are shown as thick colored lines and small rectangles, respectively. Enriched motifs are distinguished class 1 (gray) or class 2 (black). RNP-MaP densities are the number of RNP-MaP sites observed within a 25 nt window. All densities are on the same scale, with 19 as the maximum density achieved (in-cell). Locations of eCLIP sites for other proteins are shown at bottom. (b) RNP-MaP enriched sequence motifs (by MEME)^75^ and E-values for each tested protein, shown as position weighted matrices, for class 1 and class 2 motifs. (c) Relative expression levels of reporter RNAs containing non-XIST control, XIST E region, and other highly interactive XIST (by RNP-MaP) sequences, shown for both cytoplasmic and nuclear fractions. E region insert includes nts 11900-13100 of XIST. Nucleotide positions in XIST for other inserted protein-interactive sequences are indicated. qPCR signal from reporters were first normalized to 18S ribosomal RNA signal in each fraction, and then each reporter signal was normalized to the non-XIST control. Error bars represent standard error of the mean (n=3). All differences between E region reporter and other constructs are significant (P < 0.02) by Student’s t-test (two-sided), except for the difference between XIST E and Non-XIST nuclear fractions (P = 0.28). (d) XIST E region containing reporters localize in condensed foci (by *in situ* hybridization, ISH, white arrows) that co-occur with deformed nuclei (by DAPI), while non-XIST control reporter shows diffuse localization pattern with no nuclear deformation. White scale bars indicate distances of 5 µm.

Strikingly, the patterns of through-space RNP-MaP correlations for the RNase P and RMRP RNAs are nearly identical after alignment by structural domains (Fig. 3b). For RNase P, the strongest correlations define three hubs involving the specificity domain, the substrate cleavage site, and the Rpp20/25 dimer (which links the specificity domain and cleavage site), and these hubs are conserved in RMRP (Fig. 3c). Omission of protein complexes that comprise each hub suppresses or eliminates RNase P catalytic function *in vitro*^25^. Our RNase P and RMRP data thus identify shared protein interactions and interaction network hubs in two structurally related but sequence divergent RNAs, confirm conservation of these interaction networks between mice and humans, and highlight the ability of RNP-MaP to identify functional regions in large noncoding RNAs.

### RNP-MaP identifies conserved protein interaction networks in the XIST lncRNA

The 20-kb X-inactive specific transcript (denoted Xist in mouse, XIST in human) is a lncRNA that controls X-chromosome dosage compensation in eutherian mammals^26^. The Xist/XIST sequence is only minimally conserved between mice and humans, despite accomplishing the same functions. We used RNP-MaP to examine whether Xist/XIST protein interaction networks are conserved in the absence of sequence conservation. After in-cell crosslinking, Xist/XIST RNAs were enriched by RNA antisense pull-down^27^, and RNA-protein interactions were read out by randomly primed MaP RT (Fig. S7). Data were obtained for 97% of nucleotides in both RNAs, including over four nucleobases and in both structured and unstructured regions (Fig. S8).

Strikingly, regions with high RNP-MaP site density in Xist/XIST corresponded to the A, B, C, D, and E regions (Fig. 4a), which are each known to contain distinct repetitive sequences important for Xist localization, assembly, and chromatin silencing^28–35^. High RNP-MaP density was observed at each of these repetitive regions despite changes in copy number (human XIST contains two copies of the B region) and changes in size, relative position, and sequence (C, D, and E regions differ extensively between mice and humans)^36^. We also observed multiple additional regions of Xist/XIST, not previously defined in terms of Xist function, with strong RNP-MaP signal density that are clearly conserved between mice and humans.

We compared the RNP-MaP signal to eCLIP site density, measured in experiments in K562 cells^37^, from 30 proteins whose binding sites on XIST have been mapped and were reproducible between eCLIP replicates. We observed good agreement between RNP-MaP and eCLIP data, especially over the human XIST A and E regions (Fig. 4b). The eCLIP sites of human HNRNPK protein, whose mouse homologue mediates recruitment of Polycomb through the B and C regions of mouse Xist^29, 38^, closely mirror RNP-MaP site density across the human XIST D region and overlap human XIST B and C regions (Fig. S8), supporting a model wherein the B, C, and D regions of Xist/XIST are functionally analogous^39^. eCLIP site density tends to be lower than RNP-MaP density in multiple regions, for example between the two human XIST B regions, likely reflecting that only a subset of Xist-binding proteins have been mapped by eCLIP. Together, these data show that RNP-MaP efficiently identifies protein-binding sites critical for function in very long lncRNAs.

RNP-MaP correlation patterns in the Xist/XIST RNAs showed that protein-bound regions form higher-order interaction networks with distinct levels of interactivity, features invisible to alternative analysis strategies. Six highly networked regions were detected in human XIST RNA, and at least five of these regions are conserved in mouse Xist RNA (Fig. 4a). Low sequencing depth precluded identification of high-confidence correlations in the human XIST A region. The E region of Xist/XIST represents an extreme example in which an extended region (spanning 1-1.5 kb) forms a highly cooperative protein interaction network as evidenced by high correlation strength densities. There exist other highly protein-bound regions, such as the C region of mouse Xist and portions of Xist/XIST D region, that are highly bound but do not show correspondingly strong correlations (Fig. 4a); in these regions proteins that bind do so more independently of one another. Thus, RNP-MaP data identify distinct local patterns of RNA-protein interaction networks, detected as low and high levels of network interactivity.

### Communities of XIST-binding proteins

We next used RNP-MaP correlations to directly analyze inter-protein communication in the XIST RNP. We performed a network analysis of high-confidence eCLIP sites that are linked by RNP-MaP correlations to reveal communities of XIST-interacting proteins. We identified four communities of XIST-binding proteins by maximizing modularity of the network^40, 41^ while weighting by correlation strength (mutual information), (Fig. 5, **Table S1**). We categorized these communities based on the functions of XIST sequences to which the proteins bind, including 5′ Silencing, Compartmentalization, Splicing, and U/C communities (Fig. 5 **and** Fig. S8). The communities are distinct: correlations between proteins from different communities occur significantly fewer times than expected based on the proximity of their binding sites (**Table S2**).

Proteins in the 5′ Silencing community bind primarily to the 5′ region of XIST, which includes the silencing-critical A region^28^. Community members include factors involved in XIST processing, XIST stability, and XIST-mediated silencing - UCHL5^42^, EXOSC5^43^, HNRNPUL1^44^, and RBM15^45^, (Fig. 5). Silencing community members TARDBP and RBM22 are RNA-dependent regulators of transcription^46^ but have not yet been directly implicated in X chromosome silencing. The significant interactivity of the binding sites of the 5′ Silencing community members suggests formation of a specific coordinated RNP on XIST.

The strongest inter-protein correlations are observed between Compartmentalization community members PTBP1, MATR3, and TIA1, which bind in the XIST E region (Fig. 5 **and** Fig. S8). XIST E region is critical to maintenance of the silenced X chromosome compartment^33^. Notably, PTBP1, MATR3, and TIA1 each undergo liquid-liquid phase-transitions to form RNA granules^47–50^, a feature consistent with the formation of an XIST-mediated compartment. PTBP1 and MATR3 interact together on other RNAs as well, creating multi-valent interactions typical of RNA-protein granules^51, 52^.

The Splicing community includes components known to control splicing (U2AF2, SRSF1, TRA2A, AQR, and ILF3)^53–55^ and a chromatin modulator (GRWD1)^56^. All Splicing community proteins, except for TRA2A, bind to XIST at exon-exon junctions (Fig. S8), consistent with a function in splicing of XIST transcripts.

The smallest community (U/C) includes two HNRNPs (U and C) already known to interact with one another^57^. HNRNPs U and C do not strongly interact with other communities and interact sparsely across XIST, suggesting that the roles these proteins play in X chromosome silencing do not require significant interaction with other networks. Together, this analysis reveals how RNP-MaP can define RNP communities with distinct levels of networking (low versus high), associated with lncRNA functions.

### PTBP1, MATR3, and TIA1 recognize XIST E region via two binding modes

The XIST E region is critical for maintaining the silenced X chromosome compartment^32, 33^, is distinguished by strong inter-protein network connectivity (Fig. 4a), and includes binding sites for the proteins most strongly linked by RNP-MaP correlations: PTBP1, MATR3, and TIA1 (Fig. 5 **and** Fig. S8). To assess how each protein interfaces with the XIST E region, we used RNP-MaP to examine binding by purified recombinant PTBP1, MATR3, and TIA1 to *in vitro* transcribed RNA spanning the human XIST E region.

PTBP1, MATR3, and TIA1 each bound the XIST E region RNA with similar patterns (Fig. 6a). Both the quantity and reactivity of RNP-MaP sites increased in a protein concentration-dependent way (Fig. S9). Comparing in-cell RNP-MaP sites in the XIST E region to *in vitro* RNP-MaP sites for PTBP1, MATR3, and TIA1 revealed two patterns: (*i*) regions with similar *in vitro* and in-cell RNP-MaP signals and (*ii*) regions with high in-cell RNP-MaP signals not observed *in vitro* (Fig. 6a). In addition, a notable number of RNP-MaP sites for PTBP1, MATR3, and TIA1 observed *in vitro* were not identified by eCLIP. Thus, the full list of proteins that bind to the XIST E extends beyond those currently mapped by eCLIP, and the CLIP data do not include all XIST E sequences capable of being bound by PTBP1, MATR3, and TIA1.

Two distinct classes of 9-mer motifs were bound by each protein (Fig. 6b). The pyrimidine-rich motifs of class 1 are similar to motifs previously defined for each protein *in vitro*^58^ and by CLIP methods^59–61^, whereas the motif class 2 is uniquely identified by RNP-MaP. E-values for motifs of class 1 were not significant because of the high frequency of uridine nucleotides in the E region (36%) and the low complexity of the motifs. In contrast, the purine-rich class 2 motif was significantly enriched in RNP-MaP sites from each protein *in vitro* (Fig. 6b). Class 1 and 2 motifs were also found by analysis of RNP-MaP sites identified in cells, with enrichment scores comparable to those observed *in vitro* (Fig. 6b). Under our simplified *in vitro* binding conditions, PTBP1, MATR3, and TIA1 confer higher RNP-MaP reactivity on class 2 motifs than the class 1 motifs (Fig. S9 **and** S10). In cells, RNP-MaP densities and reactivities at class 1 and 2 motifs are more similar (Fig. S10). Thus, our data suggest that PTBP1, MATR3, and TIA1 recognize overlapping regions of the XIST E region through two distinct modes of recognition that are likely influenced by the presence of other XIST-binding proteins.

Interestingly, while RNP-MaP readily detects binding at both motif classes, class 2 motifs were not detected by eCLIP for PTBP1, MATR3, or TIA1 (Fig. 6a). This discrepancy likely reflects that eCLIP signal relies largely on uridine nucleotides to achieve crosslinking^8^ and the density of uridine nucleotides is low in the XIST E region where class 2 motifs are present (Fig. S10). Overall, RNP-MaP identifies a more comprehensive set of protein-binding sites such that 97% of eCLIP sites in XIST (of the 151 analyzed) overlap RNP-MaP sites, but only 35% of RNP-MaP sites (of 3766) exist within an eCLIP site. RNP-MaP does not rely on specific nucleotides for reactivity or enrichment by antibody binding and thus can identify more diverse protein recognition motifs, but RNP-MaP does not independently assign an interaction site to a specific protein.

### Protein-interactive E region promotes RNA focus formation in cells

PTBP1, MATR3 and TIA1 proteins are implicated in formation of RNA foci^47–50^, likely through multivalent interactions between RNAs and proteins^62^, consistent with the highly interactive protein networks we measure in the XIST E region (Fig. 4a). To examine whether protein interactions with the E region can promote foci formation, we inserted the E region into an RNA reporter and compared its expression and localization to reporters containing other highly protein-interactive XIST regions or a non-XIST sequence. Reporter RNA containing the XIST E region was expressed at significantly lower levels than RNA reporters containing other highly interactive XIST sequences (Fig. 6c). Furthermore, the E region-containing reporter, but not reporters containing other regions of XIST, formed large foci in cells (Fig. 6d). These foci, formed in the cytoplasm by reporter RNAs containing the E region, are reminiscent of nuclear XIST particles that form on chromatin during X chromosome inactivation^28^. Thus, the highly interconnected RNP network that assembles on the XIST E region, likely in concert with binding by granule-associated proteins PTBP1, MATR3, and TIA1, can organize an RNA into XIST-like foci. Our data support the role of the E region in organization of the XISTcompartment^32, 33^ and highlight the ability of RNP-MaP to discover and characterize novel functional motifs in noncoding RNAs.

## DISCUSSION

Here we demonstrate that RNP-MaP enables rapid and experimentally concise characterization of functionally important protein-RNA interaction networks: Protein binding sites are identified across an RNA with low sequence and structure biases, mutually exclusive interaction networks are distinguishable by their correlation patterns, highly interactive hubs are identified, and functionally important regions are assessed by their binding site density and interconnectivity.

RNP-MaP revealed new insights into the assembly of small RNPs U1, RNase P, and RMRP. Each RNP has three or four interaction network hubs. Intriguingly, the strongest protein network hubs in each RNA correspond to regions known to be central to RNP assembly and activity: the Sm complex assembly site in U1 and the substrate cleavage sites in RNase P and RMRP. The next strongest hubs are regions that comprise additional critical components in each RNP: the SNUPN protein binding site, which traffics the Sm-bound U1 RNA into the nucleus, and the specificity domains of RNase P and RMRP, which recruit substrates to the active sites. The unique ability of RNP-MaP to rank interaction networks by correlation strength will aid in discovery and prioritization of functional elements in noncoding RNAs.

Prior structure probing analyses of the mouse Xist RNA revealed that repeat-containing regions are structurally dynamic and accessible for protein binding, and these regions were proposed to function as “landing pads” for proteins^16^. Our RNP-MaP study now reveals that repeat-containing sequences in Xist/XIST are extensively bound by proteins and that these interactions are often highly networked. Granule-associated proteins PTBP1, MATR3, and TIA1 bind throughout the XIST E region, and the E region promoted foci formation by a heterologous reporter RNA in cells. These data suggest that protein interaction networks in the E region contribute to formation of a phase-separated particle, thus playing a role in XIST compartmentalization *in vivo*. Similar roles for Xist repeats, including in the E region, have recently been proposed based on analysis of Xist particle shape and composition^63^. Importantly, RNP-MaP directly identified hubs of conserved function in the absence of additional information and for RNAs lacking extensive sequence conservation. Our study additionally mapped many new highly-networked hubs that merit detailed further study.

There are estimated to be at least as many noncoding RNA genes as there are protein-coding genes^64^. It remains challenging to discern the overall functions of noncoding RNA transcripts and to identify function-critical regions of a long noncoding RNA. Conserved macromolecular structure implies function and, in the case of eukaryotic noncoding RNPs, structure and function are reliant on RNA-protein interaction networks^65^. The ability of RNP-MaP to identify function-critical regions of RNA and their interconnected protein networks *de novo* is likely to have a significant impact on understanding of the thousands of noncoding RNAs whose functions and protein networks are unexplored. RNP-MaP can be further applied to reveal how interaction networks form and dissociate during complex assembly in both coding and non-coding RNAs, how networks differ between cell types, and how networks change in response to stimuli.

## ACKNOWLEDGEMENTS

The work was supported by grants from the US National Science Foundation (MCB-1121024) and National Institutes of Health (R35 GM122532) to K.M.W. C.A.W. is a postdoctoral fellow of the American Cancer Society (ACS 130845-RSG-17-114-01-RMC). J.M.C. was supported by NIH grant R01 GM121806. Xist and XIST antisense probes were provided by the Guttman laboratory (CalTech), and we thank Mario Blanco for his initial support in their application. XIST eCLIP data from published work were provided upon request by the Yeo laboratory (UCSD), and we thank Gene Yeo (UCSD), Meredith Corley (UCSD), and Daniel Sprague (UNC) for support in formatting these data for integration into this work and for helpful comments on the project.

## AUTHOR CONTRIBUTIONS

C.A.W. and P.B.J. conducted experiments, and C.A.W., A.M.M. and K.M.W. analyzed data. C.A.W., J.M.C., and K.M.W. designed and interpreted experiments. The manuscript was written by C.A.W. and K.M.W. with input from all authors.

## DATA AVAILABILITY

Raw and processed sequencing datasets analyzed in this report will be made available upon reasonable request and deposited in the Gene Expression Omnibus database (in process).

## CODE AVAILABILITY

ShapeMapper2, deltaSHAPE, SuperFold, and RingMapper software used for analysis are available here (http://weeks.chem.unc.edu/software.html) and here (https://github.com/Weeks-UNC). MEME, VARNA, PyMol, and Gephi are freely open source software.

## METHODS

### Cell culture

Adherent mammalian cells used in chemical probing experiments, either SM33^27^ or HEK293 cells, were grown to 80-90% confluency in either 6-well plates (for targeted priming) or 10-cm dishes (for RNA antisense pulldown). HEK293 cells were cultured in DMEM with 10% FBS. SM33 cells were cultured in embryonic stem cell media [DMEM high glucose with sodium pyruvate, 15 % FBS, 0.1 mM non-essential amino acids (Gibco), 2 mM L-glutamine, 0.1 mM β-mercaptoethanol,1000 U/mL leukemia inhibitory factor (ESGRO, Millipore Sigma)]. Cultures were grown with 100 U/mL penicillin and 100 µg/mL streptomycin. To induce expression of the Xist RNA, SM33 cells were supplemented with 2 µg/mL doxycycline 16 hours before treatment. For all experiments when performing biological replicates, chemical probing and sequencing library preparation were performed on distinct populations of cells on different days.

### In-cell crosslinking with SDA

For 6-well plates, cells were washed once in 1 mL PBS, then covered with 900 µL PBS. To these cells, 100 µL of 100 mM SDA (NHS-diazirine, succinimidyl 4,4’-azipentanoate, Thermo Fisher) in DMSO was added with concurrent manual mixing. For controls, 100 µL of neat DMSO was added. Cells were treated with SDA for 10 minutes in the dark at 37 °C, then excess SDA was quenched with a 1/9 × volume of 1 M Tris-HCl, pH 8.0 (111 µL). For SM33 cells, which remained adherent during treatment, quenching was performed for 5 minutes in the dark at 37 °C. For HEK293 cells, which detached upon treatment, cells were pelleted at 1000 ×g for 3 minutes immediately after addition of quencher. Cells were washed once with PBS (and pelleted again if not adherent) and then resuspended in 400 µL of PBS in a well of a 6-well plate. To crosslink labeled proteins to RNAs, SDA-treated and untreated cells were placed on ice and exposed to 3 J/cm^2^ of 365 nm wavelength ultraviolet light (about 9 minutes in a UVP CL1000 equipped with five 8-Watt F8T5 black lights) at a distance of 4 inches from the light source. When the amount of SDA used for treatment, the amount of UV light exposure, or the compound used for crosslinking was varied no other changes were made to the procedure. When performing crosslinking procedure on cells grown in 10 cm dishes, reagent volumes used were multiplied by a factor of five relative to the 6-well procedure.

### Cellular fractionation and proteinase K lysis of SDA-treated cells

Crosslinked cells were pelleted at 1500 ×g for 5 minutes at 4 °C, washed once in cold PBS and pelleted again, and resuspended in cytoplasmic lysis buffer [10 mM KCl, 1.5 mM MgCl_2_, 20 mM Tris-HCl (pH 8.0), 1 mM DTT, 0.1% Triton X-100]. Cells were lysed for 10 minutes at 4 °C with agitation. Nuclei were pelleted at 1500 ×g for 5 minutes at 4 °C, and cytoplasmic lysates were separated into new tubes. Nuclei were washed once in low-salt solution [10 mM KCl, 1.5 mM MgCl_2_, 20 mM Tris-HCl (pH 8.0), 1 mM DTT], incubated with agitation at 4 °C for 2 minutes, pelleted again, and then resuspended in proteinase K lysis buffer [40 mM Tris-HCl (pH 8.0), 200 mM NaCl, 20 mM EDTA, 1.5% SDS, 0.5 mg/mL proteinase K]. Components were added to cytoplasmic lysates to adjust to proteinase K lysis buffer concentrations. For samples from 6-well plates, 500 µL of cytoplasmic lysis buffer and 500 µL of proteinase K lysis buffer were used; 2.5 mL of each were used for 10-cm dish samples. Nuclear and cytoplasmic fractions were incubated for 2 hours at 37 °C with intermittent mixing. Nucleic acid was recovered through two extractions with 1 volume of 25:24:1 phenol:chloroform:isoamyl alcohol (PCA) and two extractions with 1 volume of chloroform.

### Control SDA treatment of protein-free RNA extracted from cells

SM33 cells in 10-cm dishes were washed once in 5 mL PBS, then lysed in 2.5 mL proteinase K lysis buffer at 23 °C for 45 minutes. Nucleic acid was recovered through two extractions with 1 volume of PCA and two extractions with 1 volume of chloroform, and the resulting solution was buffer exchanged into PBS (PD-10 columns, GE Healthcare). The nucleic acid solution was incubated at 37 °C for 20 minutes before splitting into two equal volume portions (1.75 mL each). To one portion, a 1/9 volume of 100 mM SDA in DMSO was added, and a 1/9 volume of neat DMSO was added to the other. Each sample was incubated at 37 °C for 10 minutes in the dark. Each sample was spread evenly over a new 10-cm dish, placed on ice, and exposed to 3 J/cm^2^ of 365 nm wavelength ultraviolet light.

### In-cell treatment with 5NIA SHAPE reagent

SM33 mouse embryonic stem cells were grown in 6-well plates. In-cell 5NIA treatment proceeded as described^66^. Cells were washed once in PBS, then covered with 900 µL serum-free embryonic stem cell media. To these cells, 100 µL of 250 mM 5-nitroisatoic anhydride (5NIA, Astatech) in anhydrous DMSO was added with concurrent manual mixing. For controls, 100 µL of neat DMSO was added instead. Cells were treated with 5NIA for 10 minutes at 37 °C, cells were washed once with 1 mL of PBS, then RNA was harvested from cells with TRIzol (Invitrogen) according to manufacturer’s specifications.

### 5NIA treatment of cell-extracted RNA

SM33 cells on 10-cm dishes were washed once in ice-cold PBS and resuspended in 2.5 mL ice-cold lysis buffer [40 mM Tris-HCl (pH 8.0), 25 mM NaCl, 6 mM MgCl_2_, 1 mM CaCl_2_, 256 mM sucrose, 0.5% Triton X-100, 1000 Units/mL RNasin (Promega), 450 Units/mL DNase I (Roche)]. Cells were lysed for 5 minutes at 4 °C with agitation. Nuclei were pelleted at 1500 ×g for 5 minutes at 4 °C, resuspended in 2.5 mL of proteinase K digestion buffer, and incubated for 45 minutes at 23 °C with agitation. RNA was extracted twice with one volume of PCA that had been pre-equilibrated with 1.1× folding buffer [111 mM HEPES (pH 8.0), 111 mM NaCl, 5.55 mM MgCl_2_], followed by two extractions with one volume of chloroform. RNA was buffer exchanged into 1.1× folding buffer over a desalting column (PD-10, GE Healthcare). After incubating for 20 minutes at 37 °C, RNA solution was split into two equal portions: One was added to a 1/9 volume of 250 mM 5NIA in DMSO, and the other was added to a 1/9 volume of neat DMSO. Both portions were incubated for 10 minutes at 37 °C.

### In-cell crosslinking with photo-lysine

HEK293 cells in 6-well plates at ∼60% confluency were washed once with PBS and then cultured for 16 additional hours in media with either 2 mM natural lysine or 2 mM photo-lysine (Medchem Express). Cells were then crosslinked on ice with 10 J/cm^2^ of 365 nm wavelength UV light. Cells were washed once in PBS, pelleted at 1500 ×g, and resuspended in proteinase K lysis buffer.

Proteins were digested for 2 hours at 37 °C. Nucleic acid was recovered through two extractions with 1 volume of PCA and two extractions with 1 volume of chloroform.

### RNA precipitation and DNase treatment

Nucleic acids, including those treated after extraction from cells and those collected by Trizol or PCA after in-cell treatments, were precipitated by addition of a 1/25 volume of 5 M NaCl and 1 volume of isopropanol, incubation for 10 minutes at 23 °C, and centrifugation at 10,000 ×g for 10 minutes. The precipitate was washed once in 75% ethanol and pelleted by centrifugation at 7500 ×g for 5 minutes. Pellets from 6-well plates were resuspended in 50 µL of 1× DNase buffer and incubated with 2 units of DNase (TURBO, Thermo Fisher) at 37 °C for 1 hour. After the first incubation, 2 more units of TURBO DNase were added, and samples were incubated at 37 °C for 1 hour. Volumes were doubled for samples derived from 10-cm dishes. RNA was purified with Mag-Bind TotalPure NGS SPRI beads (Omega Bio-tek): A 1.8× volume of beads was added to DNase reactions and incubated 23 °C for 5 minutes followed by magnetic separation for 2 minutes. The solution was discarded, and beads were washed three times with 70% ethanol. RNA was eluted into 30 µL of nuclease-free water.

### Antisense-mediated purification of Xist and XIST

In 50 µL of nuclease-free water, 10 µg of total nuclear RNA (from SM33 or HEK293 cells) was heated at 70 °C for 5 minutes and then immediately placed on ice for 2 minutes. To the RNA, 100 µL of 1.5× hybridization buffer [15 mM Tris-HCl (pH 7.0), 7.5 mM EDTA, 750 mM LiCl, 0.15% Triton X-100, 6 M urea], prewarmed to 55 °C, was added. RNA was pre-cleared for 15 minutes at 55 °C with 15 µL of streptavidin-conjugated magnetic beads (Dynabeads MyOne Streptavidin C1, Thermo Fisher) that were pre-washed and resuspended in 1× hybridization buffer. After magnetic separation, the pre-cleared supernatant was retained. Biotinylated antisense RNA capture probes^27^ (Guttman laboratory Caltech), specific to either mouse Xist or human XIST, were heated at 70 °C for 5 minutes, cooled on ice for 2 minutes, then diluted in 1× hybridization buffer. Each pre-cleared RNA sample was mixed with 72 ng of capture probes, and mixtures were incubated at 55 °C for 80 minutes with shaking. After probe hybridization, 30 µL of streptavidin magnetic beads, pre-washed and resuspended in 1× hybridization buffer, were added to RNA-probe mixtures, and incubation was continued at 55 °C with shaking for 20 minutes. Beads were captured by magnetic separation and washed twice with 200 µL of 1× hybridization buffer for 5 minutes each at 55 °C. Beads were resuspended in 60 µL of NLS elution buffer [20 mM Tris-HCl, pH 8.0, 10 mM EDTA, 2% N-lauroylsarcosine, 10 mM TCEP]. RNA was eluted from beads with three heating-cooling cycles where the temperature was ramped down from 95 °C to 4 °C and up to 95 °C in 1.5-minutes cycles. Beads were captured and RNA eluates saved. The same beads were then resuspended in 40 µL of NLS elution buffer and the elution procedure was repeated; the 40 µL eluate was added to the original 60 µL eluate. Captured RNA was purified (RNeasy MinElute Cleanup Kits, Qiagen). To reduce non-target RNA in the sample, RNAs were enriched again via a second capture: the procedure was identical to first capture except omitted the pre-clear step.

### In vitro SDA crosslinking of T7-transcribed XIST E region with recombinant proteins

The E region of human XIST RNA (nucleotides 11900-13100 of NCBI NR_001564.2) was transcribed from a DNA template using T7 RNA polymerase (MEGAscript, Thermo Fisher), treated with DNase I (TURBO, Thermo Fisher), and purified via denaturing polyacrylamide gel electrophoresis. Product RNA was eluted from gels in nuclease-free water for 2 hours at 23 °C and concentrated with centrifugal filters (Amicon Ultra 10K, Millipore Sigma). Before SDA crosslinking, RNA was heat denatured at 98 °C for 2 minutes, then cooled on ice for 2 minutes before being diluted to 10 nM in 200 µl of RNP crosslinking buffer [1× PBS (pH 7.4), 1 mM MgCl_2_, 1 mM DTT] containing varying concentrations of recombinant XIST-binding proteins PTBP1, MATR3, or TIA1 (HEK293 recombinant, Origene) or BSA control protein (Millipore Sigma). RNPs were allowed to assemble for 30 minutes at 23 °C. 196 µl of mixtures were added to 4 µl of 100 mM SDA (in DMSO) in wells of a 6-well plate and incubated in the dark for 15 minutes at 23 °C. RNPs were crosslinked with 3 J/cm^2^ of 365 nm wavelength UV light. To digest unbound and crosslinked proteins, reactions were adjusted to 1.5% SDS, 20 mM EDTA, and 0.5 mg/ml proteinase K and incubated for 2 hours at 37 °C. RNA was purified once with 1.8× SPRI magnetic beads, purified again over an RNeasy MinElute column (Qiagen), and eluted into 14 µl of nuclease-free water.

### MaP reverse transcription

MaP reverse transcription was performed using a revised protocol as described^66, 67^. For smaller RNA targets (U1, RNase P, RMRP), 2 pmol of gene-specific primers were mixed with 500 ng of total nuclear RNA (or unfractionated total RNA when indicated). For MaP reverse transcription of enriched Xist/XIST RNAs or in vitro crosslinked XIST E region, 7 µL of final RNA product was mixed with 200 ng of random nonamer DNA oligonucleotides. When performing MaP reverse transcription on ribosomal RNA, 3 µg of total cytoplasmic RNA was mixed with 200 ng of random nonamers. To RNA-primer mixes, 20 nmol of dNTPs (5 nmol each base) was added (10 µL total volume of RNA, primers, and dNTPs), heated to 70 °C for 5 minutes, and then immediately placed at 4 °C for 2 minutes. To this template solution, 9 µL of freshly-made 2.22× MaP buffer [111 mM Tris-HCl (pH 8.0), 167 mM KCl, 13.3 mM MnCl_2_, 22 mM DTT, 2.22 M betaine] was added, and the mixture was incubated at 25 °C for 2 minutes. After adding 200 units of SuperScript II reverse transcriptase (Thermo Fisher), reaction mixtures were incubated for 10 minutes at 25 °C, 90 minutes at 42 °C, cycled 10 times between 42 °C and 50 °C with each temperature incubation 2-minutes long, and then heated to 70 °C for 10 minutes to inactivate enzyme. Reverse transcription reactions were buffer exchanged into TE buffer [10 mM Tris-HCl (pH 8.0), 1mM EDTA] (Illustra G-50 microspin columns, GE Healthcare).

### Two-step PCR of small RNA MaP libraries

Small RNA sequencing libraries were generated using a two-step PCR strategy as described^66, 67^. Briefly, 3 µL of cDNA from the reverse transcription reaction was used as template for step 1 PCR, using 20 cycles of gene-specific PCR (Q5 hot-start polymerase, New England Biolabs): 30 s at 98 °C, 20 × [10 s at 98 °C, 30 s at gene-specific annealing temperature, 20 s at 72 °C], 2 min at 72 °C. Each set of step 1 primers contained the same added handles to prime step 2 PCR, in which Illumina adapters and multiplex indexing sequences were appended to the libraries. Step 1 PCR products were purified (SPRI beads, Mag-Bind TotalPure NGS, Omega Bio-tek, at a 1× ratio), and 2 ng of product was used as template for step 2 PCR. Step 2 PCR involved 30 s at 98 °C, 10 × [10 s at 98 °C, 30 s at 66 °C, 20 s at 72 °C], and 2 min at 72 °C. Step 2 PCR products were purified with SPRI beads at a 0.8× ratio and eluted into 15 µL of nuclease-free water.

### Second-strand synthesis, fragmentation, and amplification of long RNA MaP libraries

For products of randomly primed MaP reverse transcription, buffer-exchanged cDNA was diluted to 68 µL with nuclease-free water. Each diluted cDNA was mixed with 8 µL of 10× Second Strand Synthesis Reaction Buffer and 4 µL Second Strand Synthesis Enzyme Mix (NEBNext, New England Biolabs), and reactions were incubated at 16 °C for 2.5 hours. The double-stranded DNA (dsDNA) products were purified with SPRI beads at a 0.8× ratio to favor longer products and exclude probe-templated products. Products were eluted into 15 µL of nuclease-free water. The dsDNA libraries were fragmented, multiplex indexed, and PCR amplified. To fragment libraries from total cytoplasmic RNA, 5 µL of 0.2 ng/µL dsDNA was combined with 10 µL of Tagment DNA Buffer and 5 µL of Amplicon Tagment Mix (Nextera XT DNA Library Prep Kits, Illumina). Mixtures were incubated at 55 °C for 5 min, then cooled to 10 °C. As soon as the temperature reached 10 °C, 5 µL of NT Buffer (Nextera XT DNA Library Prep Kits, Illumina) was added to neutralize the reaction, which was then incubated at 23 °C for 5 min. The entire reaction volume was used as a template for PCR with 15 µL of Nextera PCR Master Mix and 5 µL each of forward and reverse indexing primers (Nextera XT DNA Library Prep Kits, Illumina): 72 °C for 3 min, 95 °C for 30 s, 12 × [95 °C for 10 s, 55 °C for 30 s, 72 °C for 30 s], and 72 °C for 5 minutes. The final PCR products were purified with SPRI beads at a 0.65× ratio and eluted into 15 µL of nuclease-free water. For low concentration Xist and XIST capture libraries, 8 µL of capture product was fragmented with only 2 µL of Amplicon Tagment Mix, the concentration of index primers was halved during PCR, and PCR cycles were increased to 20.

### Sequencing of MaP libraries

Size distributions and purities of amplicon and randomly primed libraries were verified (2100 Bioanalyzer, Agilent). Step 2 amplicon libraries (about 120 amol of each) were sequenced on a MiSeq instrument (Illumina) with 2 × 150 or 2 × 250 paired-end sequencing, depending on the length of the RNA target. Libraries derived from total cytoplasmic RNA were sequenced with 2 × 300 paired-end sequencing on a MiSeq instrument, combining reads from multiple runs until desired ribosomal RNA sequencing depth was achieved. Xist and XIST capture libraries were sequenced to desired depth via a combination of 2 × 300 paired-end runs on a MiSeq and 2 × 150 paired-end runs on a NextSeq 500 instrument.

### Mutation counting and SHAPE profile generation with ShapeMapper 2 software

FASTQ files from sequencing runs, with the exception of capture libraries, were directly input into the ShapeMapper 2 software^68^ for read alignment and mutation counting. To ensure mutation rates were not affected by reduced fidelity at reverse transcription initiation sites, reads from capture libraries were trimmed by 14 nucleotides (primer length + 5 nts) after adapter sequences on each end. To accomplish this step for amplicon libraries, target FASTA files input to ShapeMapper 2 had primer-overlapping sequences and the first 5 nucleotides transcribed in RT set to lowercase, which eliminates these positions from analysis. To expedite analysis of long RNAs like Xist/XIST, corresponding FASTQs were split into ∼10 subsets and run in multiple parallel ShapeMapper 2 instances before having their outputs recombined into single profiles. ShapeMapper 2 was run with --min-depth 5000 and --output-classified flags with all other values set to defaults. In an RNP-MaP experiment, the SDA+UV-treated samples are passed as the “modified” samples and UV-only treated samples as “unmodified” samples. The outputs “profile.txt”, “parsed.mut”, and “.map” files are required for RNP-MaP site, RNP-MaP correlation, and SHAPE analyses.

### Identification of low SHAPE, low Shannon entropy regions of Xist and XIST using SuperFold

The SuperFold analysis software^13^ was used with in-cell and cell-extracted 5NIA experimental SHAPE data from mouse Xist and human XIST to inform RNA structure modeling by RNAStructure^69^. Default parameters were used to generate base-pairing probabilities for all nucleotides (with a max pairing distance of 600 nt), Shannon entropies for each nucleotide, and minimum free energy structure models.

### ΔSHAPE of mouse RNase P and Rmrp

Normalized SHAPE reactivities for 5NIA-treated mouse RNase P and Rmrp RNAs were compared between in-cell treated samples and those treated after cell extraction using the ΔSHAPE program^15^. Default parameters were used, and the 5′ primer sequence, the 3′ primer sequence, and the first 5 nucleotides transcribed during the reverse transcription step were all masked to exclude them from analysis. Only nucleotides that passed the included Z-factor and standard score significance testing were mapped as ΔSHAPE sites.

### Post-processing of mutation frequencies into RNP-MaP reactivities

Per-nucleotide mutation frequencies (number of mutation events/effective read depth) for both crosslinked (SDA+UV-treated) and uncrosslinked (UV-treated) samples were calculated from output ShapeMapper 2 profiles. RNP-MaP “Reactivity” was computed as the ratio of nucleotide crosslinked mutation frequency to uncrosslinked mutation frequency (SDA+UV rate/UV only rate). Exceptions are in Figure 1C **and S1**, where “Reactivity” refers to the ratio with a no treatment control as the denominator (treatment rate/no treatment rate). To be designated as RNP-MaP sites, nucleotide positions had to pass three quality filters: (1) sites were required to have at least 50 more mutation events in the SDA+UV-treated sample than the UV-treated sample; (2) site reactivities had to exceed the nucleotide-dependent empirical thresholds described in the next section; and (3) nucleotide reactivities were required to achieve a Z-factor greater than zero..

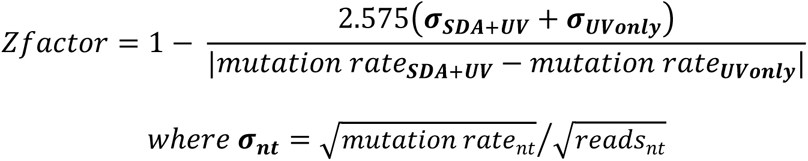

### Empirical derivation of RNP-MaP site nucleotide reactivity thresholds

Two biological replicates of RNP-MaP were performed on human U1 snRNA, RNase P RNA, and 18S and 28S rRNAs, each a part of RNA-protein complexes where atomic resolution structural data is available, enabling separation of nucleotides into two groups: those within 10 Å of protein (<10 Å) and those further than 10 Å from protein (>10 Å). For U1 snRNA, the binding site of the SNUPN protein has been mapped by crosslinking and mass spectrometry^18^, and distances between the three nucleotides surrounding the crosslink site and nearest amino acids were assumed to be less than 4 Å. For each RNA replicate, reactivities were further grouped by nucleotide identity (U, A, C, and G). The 90% reactivity value of nucleotides in a >10 Å group were set as background thresholds (BG_X>10_) and compared to the median (MED_all X_) and standard deviations (SD_all X_) of reactivities for all nucleotides included in both <10 Å and >10 Å groups to create relative threshold factors (T_X_):

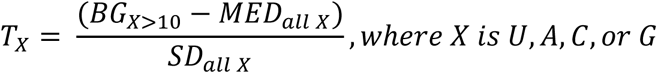

Relative threshold factors for each nucleotide from all eight replicates of the four RNAs were then averaged together, weighted by the number of nucleotides measured in each RNA, to obtain final empirically derived nucleotide relative threshold factors: 0.59 for U, 0.29 for A, 0.93 for C, and 0.78 for G. These factors represent the number of standard deviations from the median of nucleotide reactivity that must be achieved to be considered an RNP-MaP site. Factors can be applied to any RNA: to get exact nucleotide thresholds for an RNP-MaP experiment, the median reactivities for each nucleotide group (U, A, C, or G) are multiplied by their corresponding threshold factors. Factors were calculated from existing comprehensive datasets, however including more data from other RNPs or improving upon existing atomic resolution RNP structures could increase the precision of these threshold factors in the future.

### Graphical display of RNP-MaP Reactivities and crosslinking sites

Violin plots representing distributions of RNP-MaP reactivities were generated using the vioplot package in R through the web tool BoxPlotR^70^. RNP-MaP crosslinking sites were superimposed onto atomic resolution structure models using PyMol^71^. Secondary structure projection images were generated using the (VARNA) visualization applet for RNA^72^.

### RNP-MaP correlation analysis

Correlations between RNP-MaP sites were computed over 3-nucleotide windows using a previously described G-test framework (RingMapper)^19^. Windows were required to be separated by > 4 nucleotides, jointly covered by more than 10,000 sequencing reads, jointly co-mutated >50 times, and have background mutation rates below 6% (Fig. S2). Pairs of windows exhibiting G-test statistics > 20 (P < 10^-5^) in the SDA+UV treated sample and G < 10.83 (P > 0.001) in the UV-only sample were determined to be significantly correlated. Current technical limitations of MaP reverse transcription processivity (500-600 nucleotides) and sequencing instrument clustering (< 1000 nucleotides) limit distances of readily measured correlations to < 500 nucleotides.

### RNase P and RMRP structural alignment

Corresponding helices, loops, and intervening regions in RNase P and RMRP RNAs were separated into structural domains (Fig. 3) and separately aligned with MUSCLE^73^. Region alignments were recombined to create the final alignment (**Supplemental File 1**).

### Calculation of Xist/XIST RNP-MaP site, correlation strength, and eCLIP site densities

To calculate the RNP-MaP site density for Xist and XIST (Fig. 4a **and S8**), nucleotides whose reactivities were in the top 5% of all reactivities (U, A, C, and G nucleotides evaluated separately) were identified. Site density was defined as the number of nucleotides in a centered 51-nucleotide window that were top 5% sites. Correlation strength density was defined as the sum of the mutual information of all nucleotides within the window normalized to (i.e., divided by) the read depth of the central nucleotide. We selected high confidence eCLIP sites^37^ (**Supplemental File 2**) within XIST that passed Irreproducible Discovery Rate thresholds (see www.encodeproject.org/eclip), meaning that sites were defined by signal peaks of similar amplitude in both eCLIP replicates. The eCLIP site density (Fig. 4b) was defined as the total number of observed eCLIP sites within the 51-nucleotide window. For densities in Fig. 6, Fig. S9, and Fig. S10, all RNP-MaP sites were included (because signal only comes from a single protein) and centered nucleotide windows were shortened to 25 long (since the E region is only 1200 nts).

### Identification of conserved sequence regions between mouse Xist and human XIST

To identify and rank areas of significant conservation between mouse Xist (NCBI NR_001463.3) and human XIST (NCBI NR_001564.2) RNA sequences, we performed a local alignment (BLASTn^74^) and retained all segments with E-values above 0 and with lengths > 100 nucleotides and ranked these segments by alignment bitscore.

### Network analysis of XIST RNP-MaP correlation-linked eCLIP sites

To create a list of protein-linking correlations, we first counted the number of times nucleotides within our high confidence eCLIP sites were correlated in RNP-MaP data (links) and measured the summed total of mutual information within links. To ensure that links were not simply a product of eCLIP site number and proximity or the average length and density of RNP-MaP correlations, we randomly shuffled the location of RNP-MaP correlations and counted links achieved between protein pairs iteratively 2000 times, generating p-values for each protein pairing based on the number of links between them and the strength of those links (**Table S1**). Resulting links between protein pairs with p-values less than 0.05 were then used as edges connecting nodes (proteins) on a network map (Fig. 5). A maximum modularity of the network (0.419), weighted by the strengths of included links (mutual information) and without changing resolution, was calculated using Gephi^41^, and node sizes were adjusted manually to convey indicated relationships.

### Identification of RNP-MaP enriched motifs in the XIST E region

MEME^75^ was used to identify motifs enriched by RNP-MaP in vitro. We first expanded each RNP-MaP nucleotide into 9-mer sites (extending by 4 on either side). Overlapping sites were combined iteratively until no overlapping sequences remained. Combined sites were used as MEME input in classic mode using a 0-order model of sequences, allowing for any number of motif repetitions in each sequence, and explicitly looking for five 9-mer motifs. The top two motifs were retained for each in vitro experiment, and locations of all matching motifs in XIST E region were found using FIMO^76^ with a p-value threshold of 10^-3^. For the in-cell experiment, only sites from the E region were considered in MEME, and the first and fourth most significant motif (class 2 and class 1, respectively) were included in Fig. 6**, S9, and S10**.

### XIST RNA reporter plasmid design

To create XIST region-containing reporters, we used inverse PCR and re-ligation to insert a multiple cloning site into the 3′ end of the pNL 3.2.CMV vector (Promega) between XbaI and FseI sites and to add the second intron from human HBA1 at nucleotide position 196 in the nanoluciferase coding region (native Xist/XIST is spliced in its central region). Plasmids with varying regions of XIST were subcloned into the XhoI and KpnI restriction sites, and a control plasmid was generated through inverse PCR (**Table S2**).

### Plasmid transfection and purification of XIST reporter RNA for qPCR

HEK293 cells were plated at 100,000 cells/mL in 2-mL volumes per well of 6-well plates and then cultured for 24 hours at 37 °C. Each well was then transfected with a mixture of 0.6 µg of reporter plasmid and 1.8 µL of FuGENE 6 transfection reagent (Promega) in 200 µL of serum-free DMEM, and cells were cultured for an additional 48 hours at 37 °C. Transfected cells were pelleted at 1500 ×g for 5 minutes at 4 °C, washed once in cold PBS and pelleted again, and resuspended in 500 µL cytoplasmic lysis buffer. Cells were lysed for 10 minutes at 4 °C with agitation. Nuclei were pelleted at 1500 ×g for 5 minutes at 4 °C, and cytoplasmic lysates were separated into new tubes. Nuclei were washed once in low-salt solution, incubated with agitation at 4 °C for 2 minutes, pelleted again, and then resuspended in 500 µL proteinase K lysis buffer. NaCl, EDTA, SDS, and proteinase K were added to cytoplasmic lysates up to proteinase K lysis buffer concentrations. Nuclear and cytoplasmic fractions were incubated for 2 hours at 37 °C with intermittent mixing. Nucleic acid was recovered through two extractions with 1 volume of 25:24:1 PCA, two extractions with 1 volume of chloroform, and precipitation with 1/25 volume of NaCl and 1 volume of isopropanol.

### qPCR of reporter RNAs from nuclear and cytoplasmic fractions

Equal quantities of RNA (30 ng) from each nuclear and cytoplasmic fraction were used generate first-strand cDNA in random hexamer-primed reverse transcription reactions using SuperScript II (Thermo Fisher). Triplicate mixtures of 2.5 µL of template cDNA, 2.5 µL of 2 µM primers, and 12.5 µL of Maxima SYBR Green qPCR Master Mix (Thermo Fisher) in 25 µL total reactions were prepared for each sample. Reaction mixtures from matched no-reverse transcriptase controls were also prepared. Primer sets specific to the reporter gene and 18S ribosomal RNA (as a normalization control) were used for each sample fraction. qPCR was performed on a QuantStudio 6 Flex Real-Time PCR System (Thermo Fisher) with steps of 5 min at 95 °C, 40 cycles of [15 s at 95 °C, 30 s at 65 °C, 40 s at 72 °C], and a melt curve to confirm single major products. Fluorescence readings were taken at elongation steps (72 °C). All specific signals were observed more than 8 cycle thresholds earlier (256-fold more signal) than no-reverse transcriptase controls. Signals were averaged across triplicate qPCRs, normalized to 18S rRNA signal, then normalized to control reporter signals.

### RNA FISH of XIST RNA reporters

Custom RNA FISH probes antisense to the nanoluciferase mRNA and labeled with Quasar 570 Dye were ordered from LGC Biosearch with design parameters of masking level 5, oligo length 19, and minimum spacing length 2). HEK293 cells were plated at a concentration of 10,000 cells/mL in wells of a 12-well plate, with each well containing a poly-L-lysine coated #1.5 coverslip (Neuvitro). After 24 hours of growth, 50 µl of transfection mix [20 µl of 15 ng/µl reporter plasmid, 0.9 µl of Fugene 6 (Promega), and 79 µl of DMEM] was added to cells. At 48 hours post-transfection, each well was washed once with 1 ml PBS, fixed with 1 ml of 3.7% formaldehyde in PBS for 10 minutes at room temperature, then washed twice with PBS. Cells were permeabilized with 1 ml of 70% ethanol overnight at 4 °C. After removing ethanol, cells were incubated in wash buffer 1 [20% Wash Buffer A (LGC Biosearch), 10% formamide] for 5 minutes at room temperature, and then coverslips were transferred to a humidified chamber with cells facing down onto 100 µl of hybridization buffer [90% Stellaris RNA FISH Hybridization Buffer (LGC Biosearch), 10% formamide, 125 nM antisense probes]. After overnight incubation in the dark at 37 °C, coverslips were transferred into 12-well dishes and incubated in 1 ml of wash buffer 1 for 30 minutes at 37 °C in the dark, then counterstained for 30 minutes at 37 °C with 5 ng/µl DAPI in 1 ml of wash buffer 1. Coverslips were washed a final time in 1 ml of Wash Buffer B (LGC Biosearch) before being mounted onto a microscope slide with 12 µl of Vectashield Mounting Medium (Vector Laboratories) and sealed. RNA FISH z-stack images were captured using a 100×/1.3 oil objective on an Olympus IX81 Microscope and were deconvoluted using the AutoQuant X software. Z-stacks were collapsed into maximum intensity projections using the Fiji software.

---

**Figure S1.**
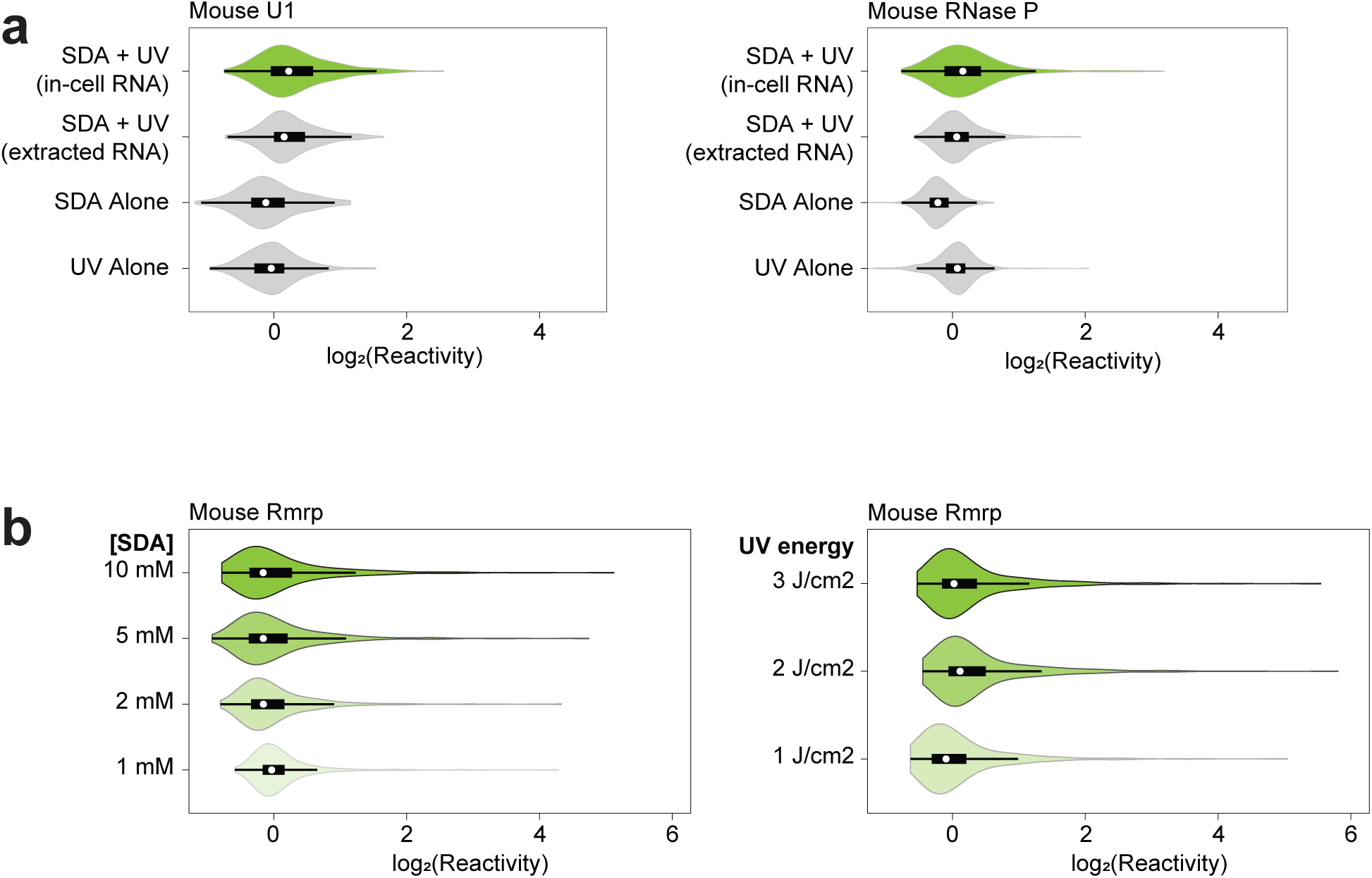
RNP-MaP signal in living cells is dependent on SDA and UV. (a) Violin plots of log_2_(reactivity) for U1 and RNase P RNA for reactions in SM33 cells, with RNA extracted from cells, and in cells without SDA, and without UV. (b) Violin plots of log_2_(reactivity) of Rmrp in SM33 cells as functions of SDA and UV doses. When SDA concentration was varied, UV dose was 3 J/cm^2^. When UV energy was varied, SDA concentration was 10 mM. For violin plots, white circles indicate medians, box limits indicate the first and third quartiles, whiskers extend 1.5 times the interquartile range, and smoothed polygons show data density estimates and extend to extreme values.

**Figure S2.**
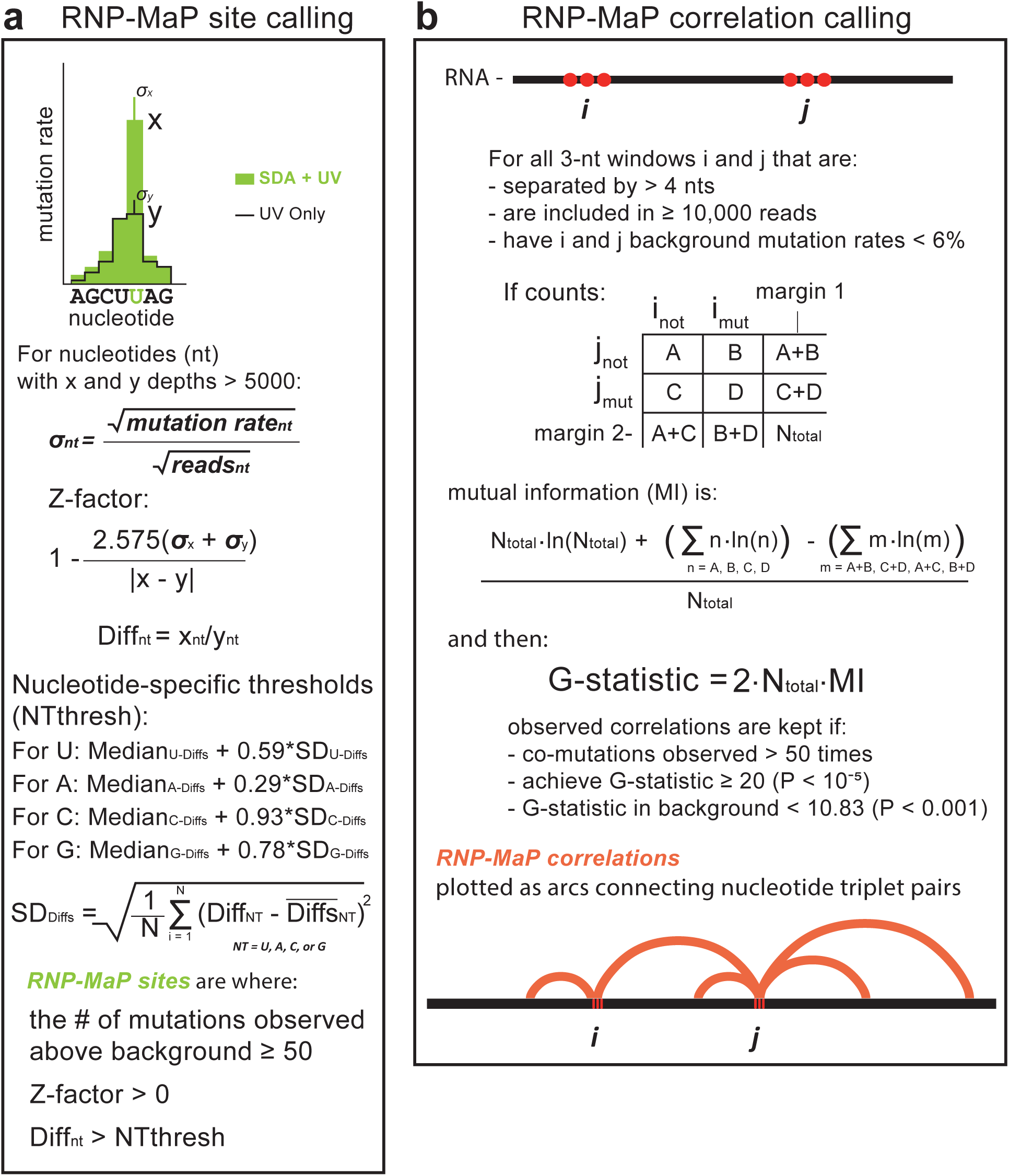
RNP-MaP site and correlation determination. Scheme outlining determination of (a) RNP-MaP sites and (b) correlations. For (a), depth threshold and mutation count above background threshold were set to eliminate noise at signal margins. A Z-factor above 0 is achieved only when 99% confidence intervals of UV only and SDA + UV rates do not overlap. To limit false positives, nucleotide-specific thresholds were set at the levels of highest reactivity among nucleotides distant from proteins in RNPs of known structure (U1, RNase P, 18S, and 28S), expressed as the number of standard deviations from median reactivities of each nucleotide (detailed in methods). For (b) 3 nt windows and 4 nt separations were set to accommodate the size of some longer and more complex mutation and deletion events during MaP reverse transcription. Thresholds of 10,000 depth, 6% background mutation rate, and 50 co-mutation events limit noise and false positives.

**Figure S3.**
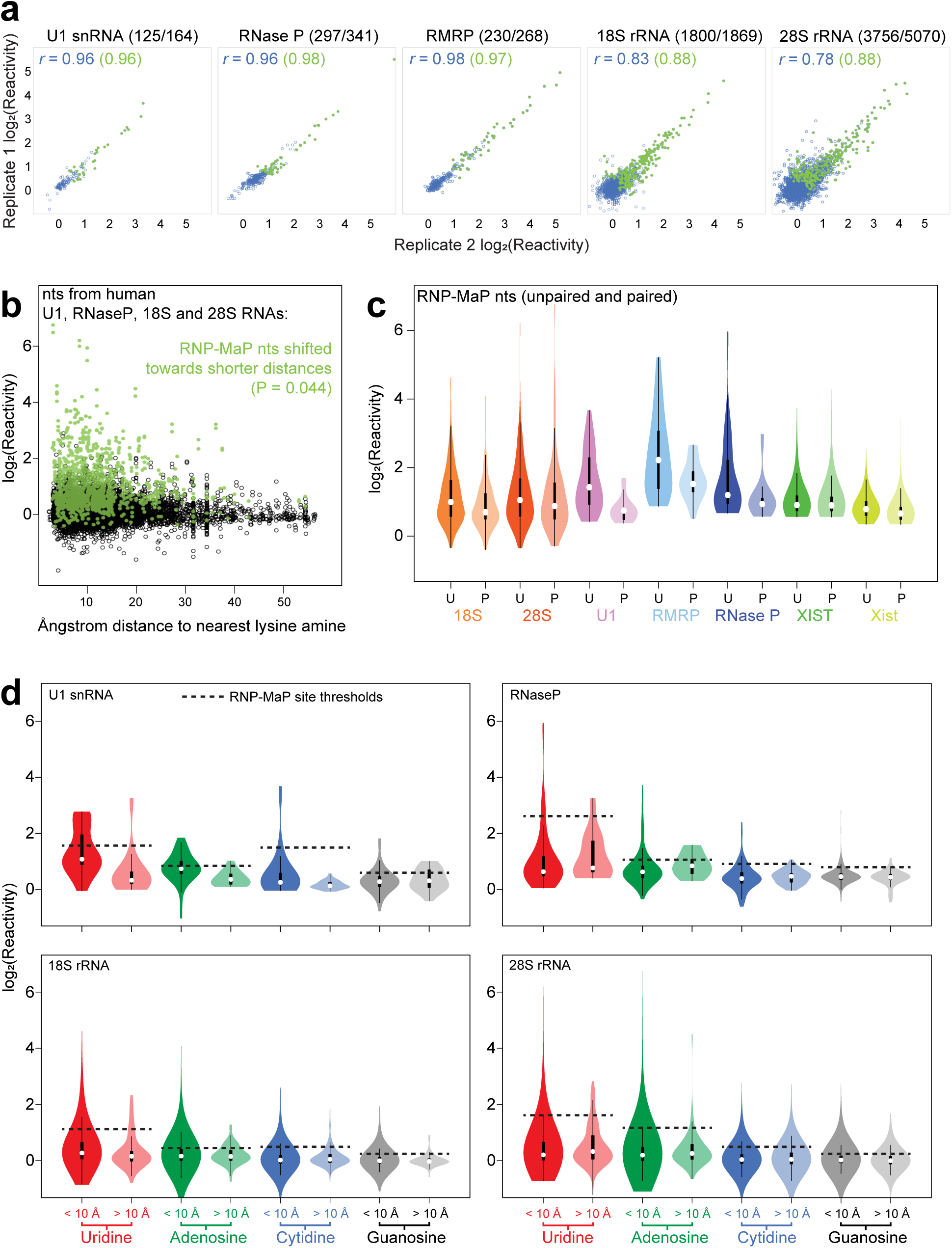
RNP-MaP reproducibly discriminates protein-bound nucleotides independent of local RNA structure and nucleotide identity. (a) Correlation (Pearson’s *r*) between log_2_(reactivity) of nucleotides from biological replicates of RNP-MaP experiments in HEK293 cells on U1, RNase P, RMRP, 18S, and 28S RNAs. is highly reproducible between biological replicates (bottom panels), especially for RNP-MaP sites. Correlations are shown for all nucleotides (blue dots) and RNP-MaP sites (green dots) for all RNAs. The number of nucleotides represented from each RNA is indicated above each graph in parentheses. (b) Nucleotide log_2_(reactivity) from RNP-MaP experiments in HEK293 cells for human U1, RNase P, 18S, and 28S RNAs are plotted against distance to protein lysine side chain amines. The significance of the difference between RNP-MaP site (green) and non-site (black) distance distributions is indicated (Mann-Whitney U-test, one-sided). (c) Violin plots of RNP-MaP site nucleotide log_2_(reactivity) from experiments in HEK293 cells (18S, 28S, U1, RNase P, RMRP, XIST RNAs) or SM33 cells (Xist RNA), grouped as unpaired nucleotides (U) and paired nucleotides (P). (d) Violin plots of nucleotide log_2_(reactivity) from RNP-MaP experiments in HEK293 cells (U1, RNase P, 18S, and 28S RNAs) grouped as protein proximal (< 10 Å) and protein-distant (> 10 Å) nucleotides and separated by nucleobase identity (Uridine, Adenosine, Cytidine, and Guanosine). To obtain nucleobase-specific thresholds (black dotted lines) for consideration as RNP-MaP sites: the standard deviations of reactivity for all of each nucleobase within an experiment are multiplied by corresponding empirically-derived threshold factors (U – 0.59, A – 0.29, C – 0.93, G – 0.78), and these products are added to the corresponding nucleobase reactivity medians. For violin plots in panels c and d, white circles indicate medians, box limits indicate the first and third quartiles, whiskers extend 1.5 times the interquartile range, and smoothed polygons show data density estimates and extend to extreme values.

**Figure S4.**
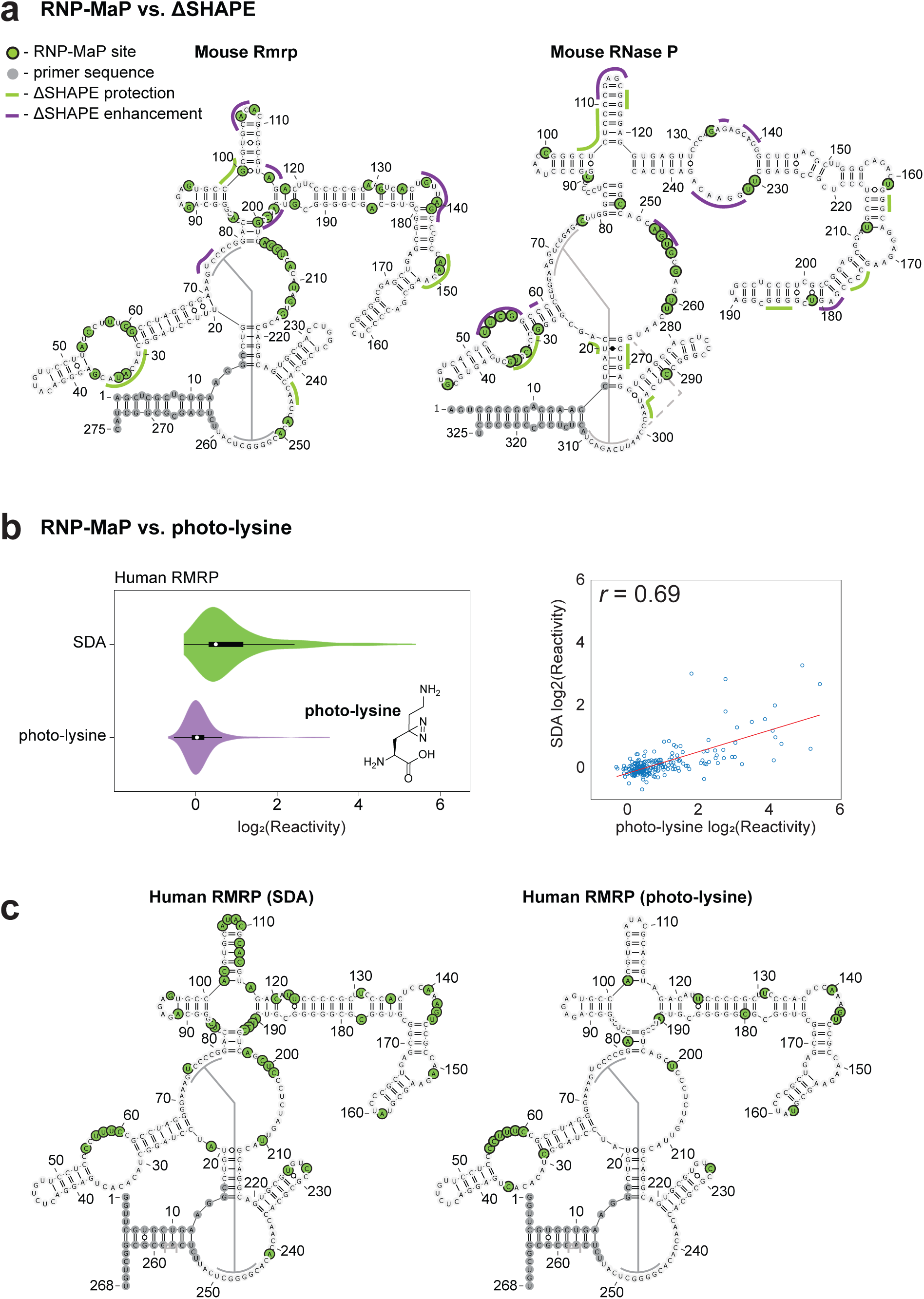
Orthogonal chemical probing and metabolic labeling approaches corroborate RNP-MaP. (a) ΔSHAPE identifies changes in SHAPE reactivity between RNA in cells and in solution, identifying regions of RNA that interact with cellular components, including proteins^15^. Overlap of RNP-MaP and ΔSHAPE sites are shown on secondary structure diagrams for mouse Rmrp and RNase P RNAs. Of 65 RNP-MaP sites (green shading) in Rmrp and RNase P RNAs, 22 (34%) are ΔSHAPE sites (lines), and 52 (80%) are within five nucleotides of ΔSHAPE sites by contact distance (which includes distances across base pairs in the secondary structures). In-cell SHAPE protections (green) and enhancements (purple) are labeled. Primer-binding sequences excluded from analysis are indicated, and solid grey and dotted grey lines indicate pseudoknot helices and predicted tertiary base-pair (inferred from human RNase P structure), respectively. (b) Metabolic labeling by photo-lysine incorporates diazirines into native lysine side chains, and these residues can be photo-activated, like SDA, with 365 nm UV light. A violin plot of log_2_(reactivity) shows that photo-lysine is less reactive with RNA than is SDA (left), as it is likely incorporated into proteins with low efficiency, has no linker, and requires a shorter crosslinking distance. White circles indicate medians, box limits indicate the first and third quartiles, whiskers extend 1.5 times the interquartile range, and smoothed polygons show data density estimates and extend to extreme values. Correlation (Pearson’s *r*) between SDA and photo-lysine reactivity (right) is shown. (c) All photo-lysine reactive regions are also reactive by SDA, as illustrated on secondary structure projections of RMRP.

**Figure S5.**
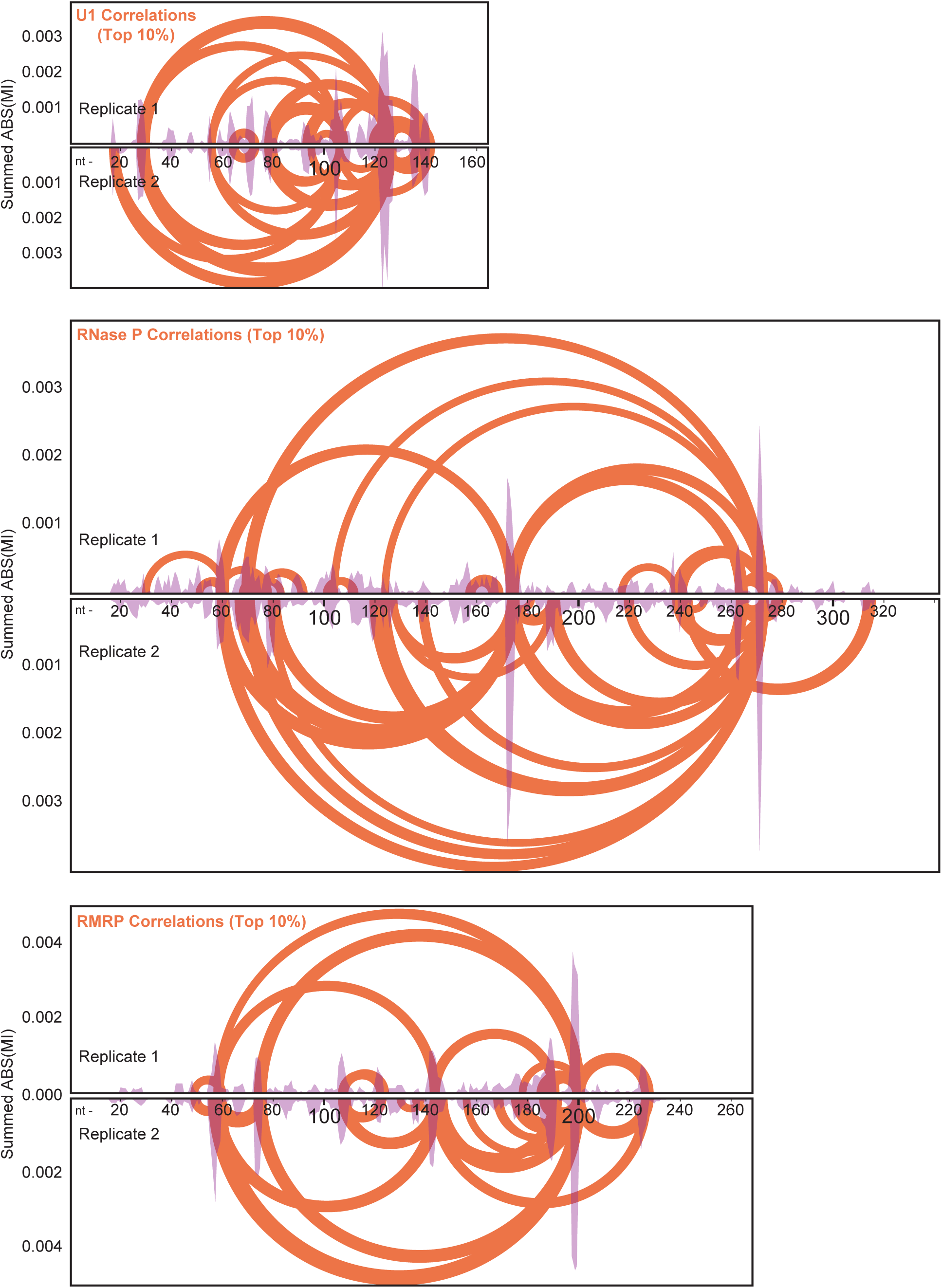
RNP-MaP interaction networks are highly reproducible. Top 10% of RNP-MaP correlations by mutual information (arcs) and the per-nucleotide summed mutual information for all RNP-MaP correlations (purple peaks) from two biological replicates in HEK293 cells are plotted by nucleotide position (nt) for human U1, RNase P, and RMRP RNAs.

**Figure S6.**
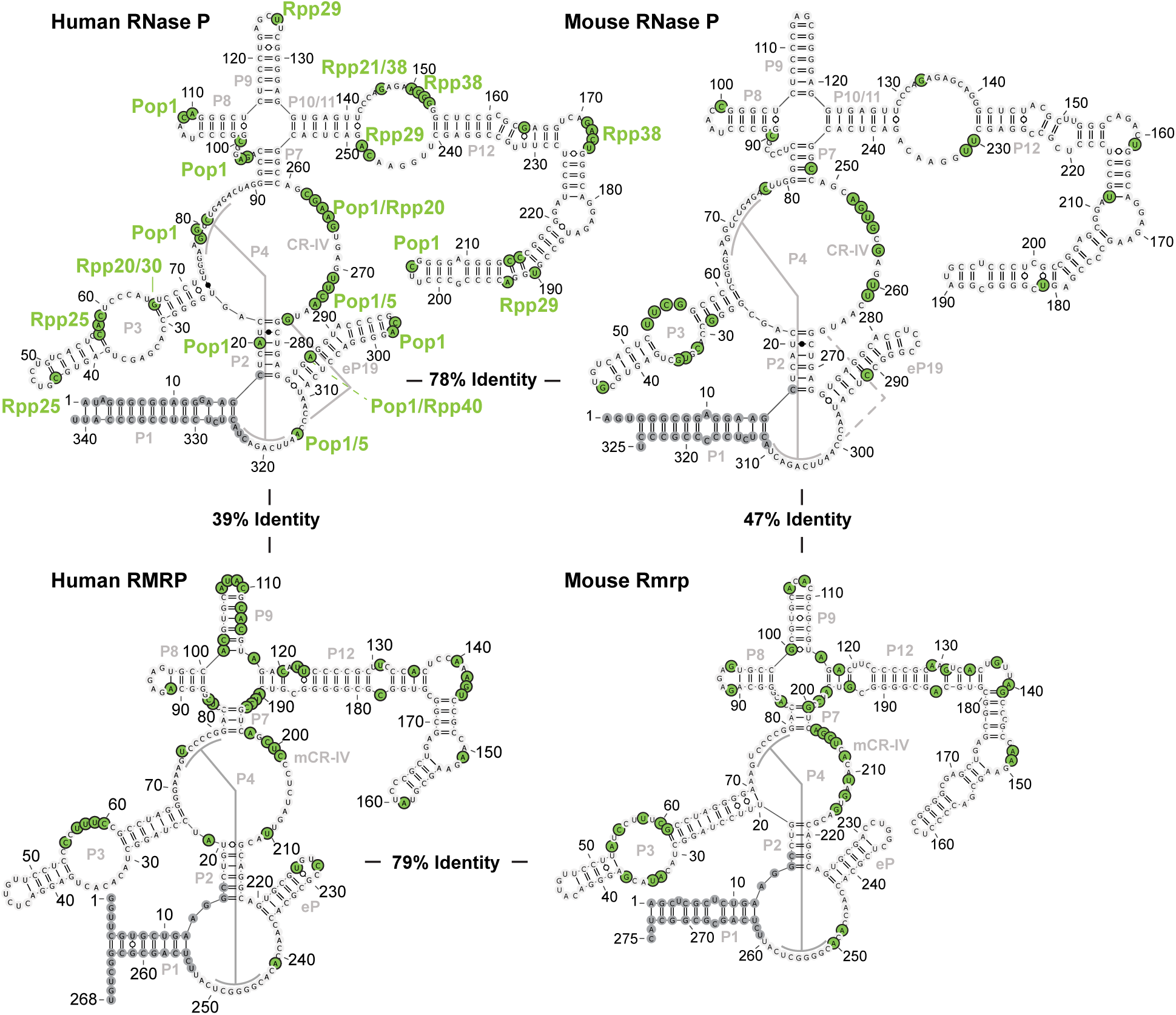
RNP-MaP sites are conserved between mouse and human RNAs. Locations of RNP-MaP sites are shared between mouse and human homologues of RNase P and RMRP, as illustrated on secondary structure projections of each RNA. Structural domains are labeled in gray, the percent sequence identities between RNAs are indicated, and gray lines denote pseudoknot helices. Proteins in close proximity to each RNP-MaP site on RNase P (based on distance to lysine amine) are labeled in green lettering. Experiments performed in SM33 or HEK293 cells.

**Figure S7.**
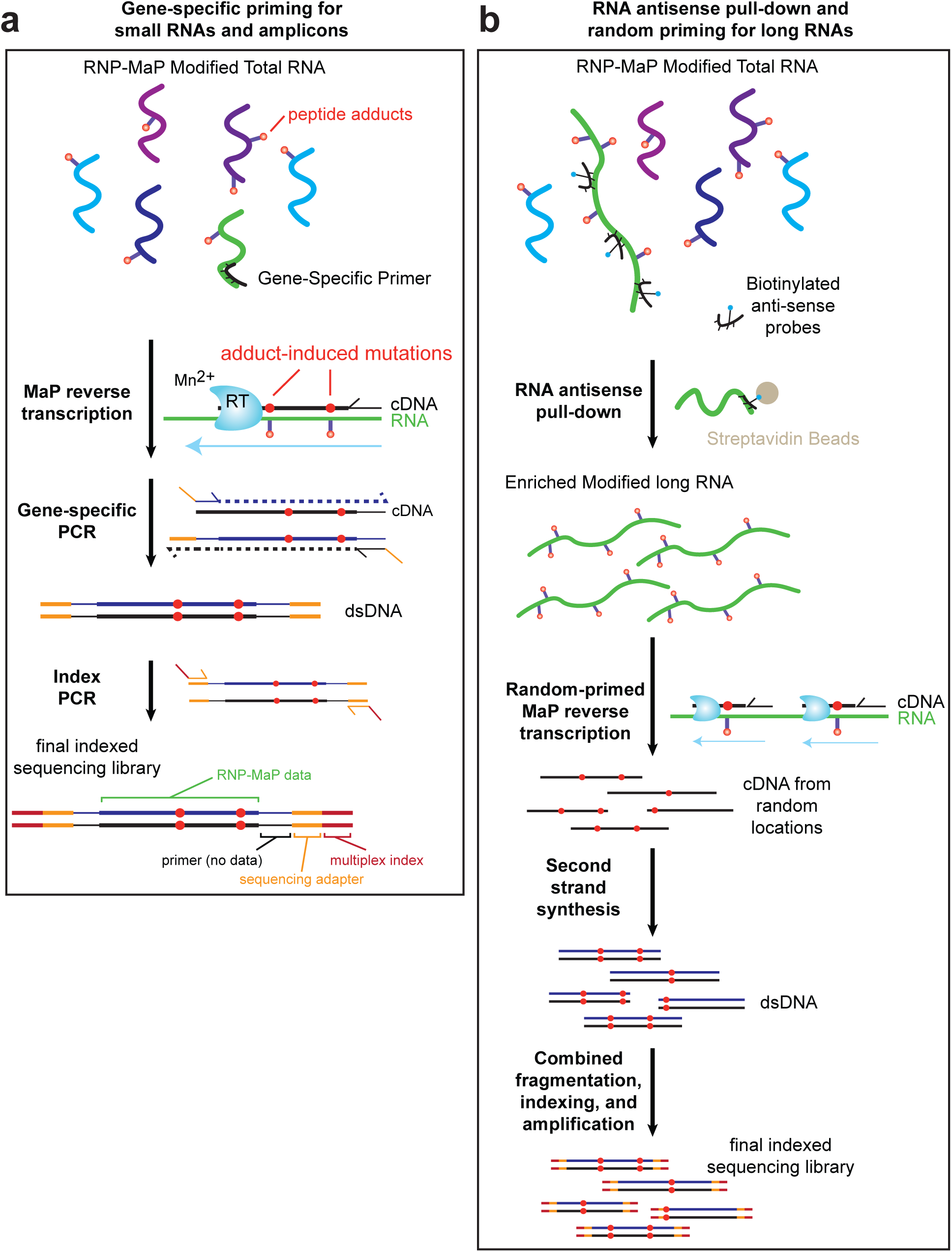
Gene-specific and random priming strategies for RNP-MaP library preparation. Schemes outlining generation of RNP-MaP sequencing libraries from (a) small RNA targets and amplicons or (b) long RNA targets are shown. Gene-specific priming is recommended for RNAs or regions of an RNA < 500 nucleotides in length, and enrichment and random priming is suggested for longer RNAs.

**Figure S8.**
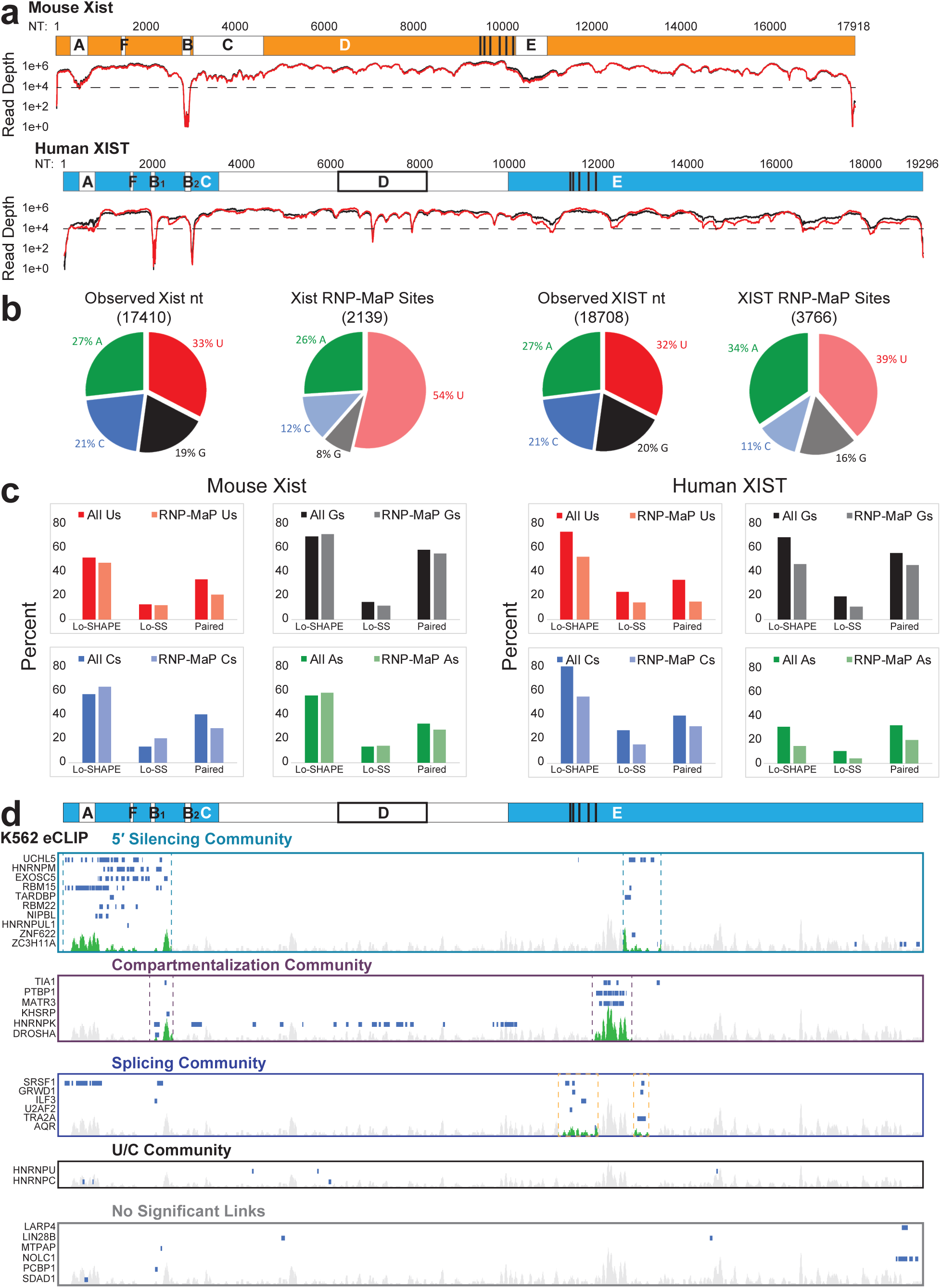
RNP-MaP captures protein interaction information across very long non-coding RNAs. (a) Almost all nucleotides in mouse Xist (orange) and human XIST (blue) were observed above 10,000 depth thresholds in RNP-MaP experiments, shown per-nucleotide. Tandem repeat sequence arrays A-F (white boxes with black letters), regions homologous to repeat regions (white letters), the D region core (white box with thick black outline), and exon-exon junctions (thick vertical black lines) are indicated on Xist and XIST. (b) RNP-MaP sites were more frequently observed at adenosine and uridine than at guanosine and cytidine (pie charts). (c) RNP-MaP signal is minimally biased toward unstructured regions in Xist and XIST, as shown by the percentages (bar graphs) of RNP-MaP site nucleotides that also have low SHAPE reactivity (Lo-SHAPE), have low SHAPE reactivity and low modeled Shannon entropy (Lo-SS), or are base paired in minimum free-energy models (paired). (d) Positions of 151 high confidence eCLIP sites for 30 XIST-binding proteins used in this analysis are shown with RNP-MaP site density (top 5% reactive sites per 51 nucleotides) aligned to the XIST sequence. Protein eCLIP sites are organized on density plots by identified community and ranked (from top to bottom) by decreasing summed mutual information of intra-community correlations.

**Figure S9.**
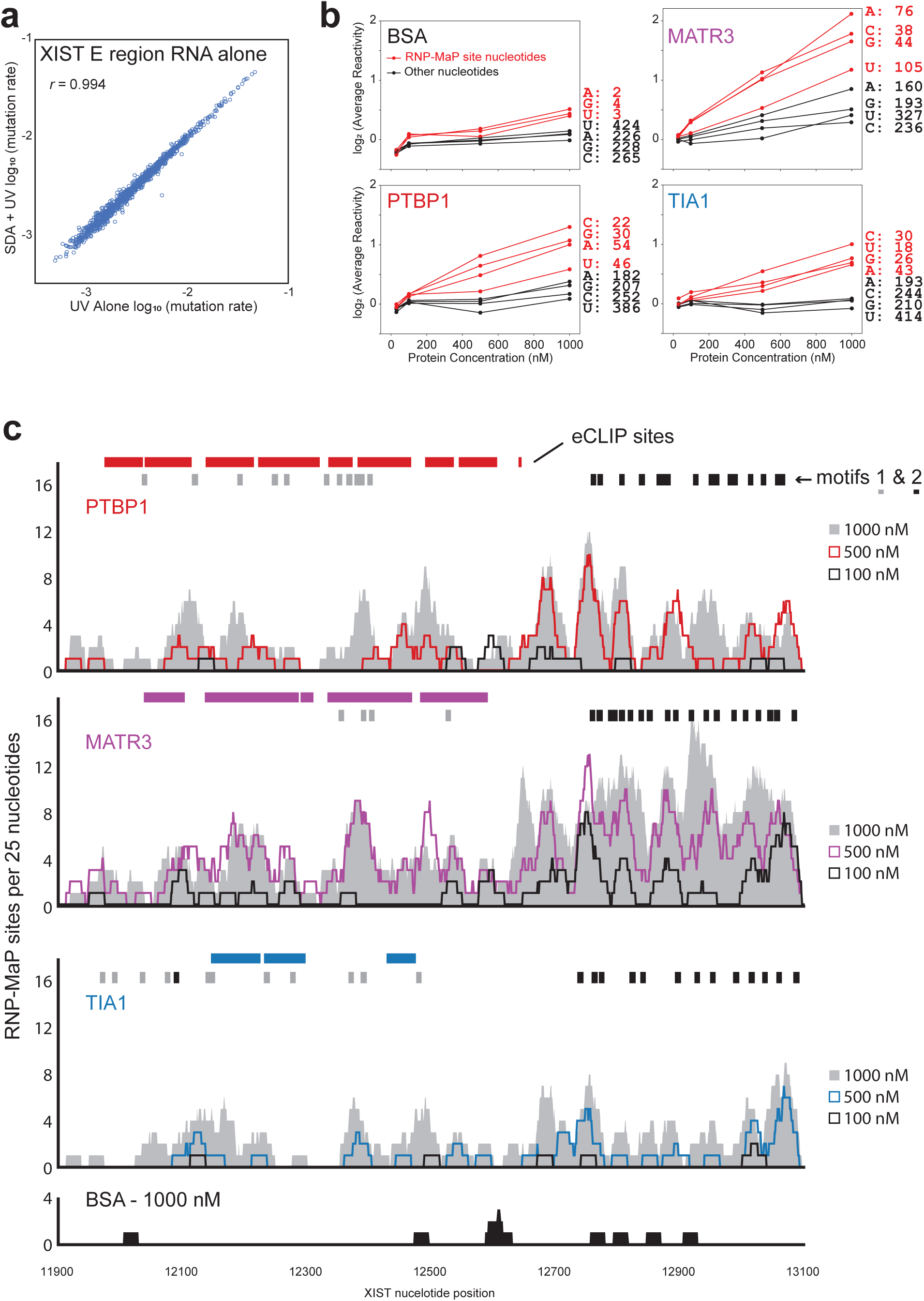
PTBP1, MATR3, and TIA1 bind the XIST E region *in vitro*. (a) Correlation of log_10_(mutation rates) of T7-transcribed XIST E region RNA with SDA and UV or UV alone in the absence of protein. MaP RT-induced mutation rates are identical (Pearson’s *r* = 0.994). (b) log_2_(reactivity) of RNP-MaP site nucleotides (red) as a function of indicated increasing protein concentration, compared to non-RNP-MaP nucleotides (black). Reactivities shown are averages for all nucleotides of each indicated nucleobase identity (right), and the number of nucleotides in each group present in each experiment is indicated. (c) Detection of RNP-MaP sites (represented as 25 nt window densities) as a function of increasing amounts of PTBP1, MATR3, TIA1, and BSA (latter is control, no effect seen). Locations of eCLIP sites and class 1 and 2 motifs for each protein are indicated on density plots.

**Figure S10.**
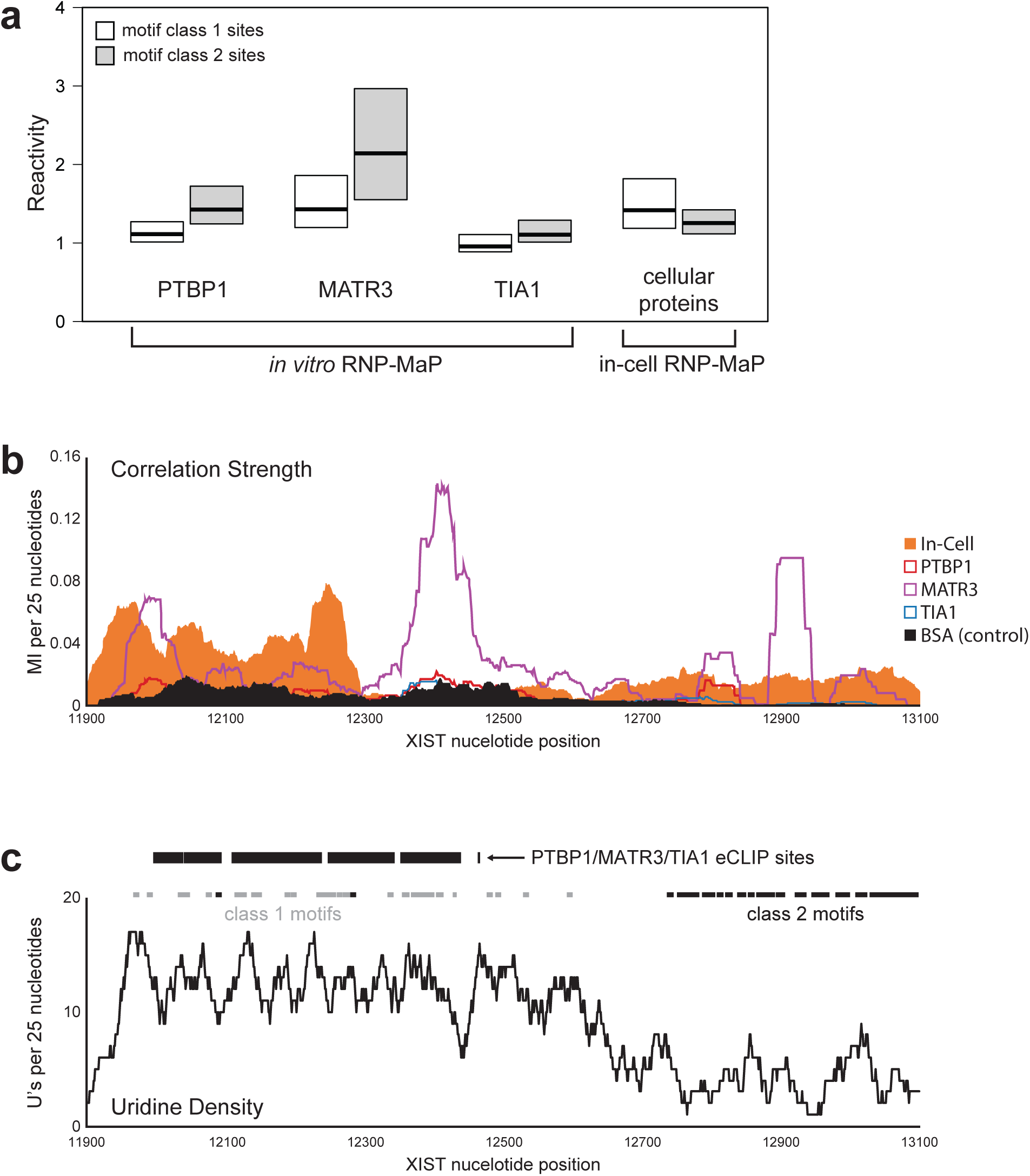
PTBP1, MATR3, and TIA1 networks observed *in vitro* do not fully recapitulate in-cell networks. (a) Recombinant PTBP1, MATR3, and TIA1 interact more strongly with motif class 2 than class 1 in XIST E region transcript *in vitro*, as shown by RNP-MaP reactivity, whereas motif class 1 is slightly more reactive in native XIST from HEK293 cells. (b) PTBP1 and TIA1 alone do not maintain significant interaction networks, as measured by mutual information (MI) density (summed over 25 nt window), and MATR3 alone *in vitro* forms networks distinct from those observed in cells. (c) PTBP1, MATR3, and TIA1 eCLIP sites are only found within the uridine-rich region containing class 1 motifs but are absent in the uridine-poor region where RNP-MaP identified class 2 motifs predominate.

**Table S1. Totals and summed mutual information (MI) of correlations linking pairwise combinations of XIST-binding protein eCLIP sites.** P-values for both attributes were generated from 2000 randomizations (without replacement) of measured correlation locations along XIST (for nucleotides with sufficient read depth only). All measured correlations are included.

**Table S2. Pairs of protein eCLIP sites significantly excluded (P-values > 0.95) from RNP-MaP data based on both total number of correlations and summed mutual information (MI) strength.** Associated network communities, total numbers of each community pairing, and the percent of pairings that were significantly excluded are also listed.

**Table S3. DNA sequences of MaP library preparation primers, plasmid construction, and qPCR primers.** Sequences inserted during plasmid construction (bold and underlined) are shown with flanking plasmid sequences included (not bold or underlined).

**Supplemental file 1. Structure-based sequence alignment of human RNase P and RMRP RNAs.**

**Supplemental file 2. List of high confidence eCLIP sites on XIST organized by protein.**

