## Supplemental Tables and Files for "RNP-MaP: In-cell analysis of protein interaction networks defines functional hubs in RNA"

| Supplemental Table S1 | | | | | |
|---|---|---|---|---|---|
| protein 1 | protein 2 | # of linking correlations | Summed MI | P (# of linking correlations) | P (Summed MI) |
| AQR | AQR | 0 | 0 | 1 | 1 |
| AQR | DROSHA | 0 | 0 | 1 | 1 |
| AQR | EXOSC5 | 0 | 0 | 1 | 1 |
| AQR | GRWD1 | 0 | 0 | 1 | 1 |
| AQR | HNRNPC | 0 | 0 | 1 | 1 |
| AQR | HNRNPK | 0 | 0 | 1 | 1 |
| AQR | HNRNPM | 0 | 0 | 1 | 1 |
| AQR | HNRNPUL1 | 0 | 0 | 1 | 1 |
| AQR | HNRNPU | 0 | 0 | 1 | 1 |
| AQR | ILF3 | 17 | 0.005105143 | 0 | 0.0015 |
| AQR | KHSRP | 0 | 0 | 1 | 1 |
| AQR | LARP4 | 0 | 0 | 1 | 1 |
| AQR | LIN28B | 0 | 0 | 1 | 1 |
| AQR | MATR3 | 17 | 0.001056889 | 0 | 0.088 |
| AQR | MTPAP | 0 | 0 | 1 | 1 |
| AQR | NIPBL | 0 | 0 | 1 | 1 |
| AQR | NOLC1 | 0 | 0 | 1 | 1 |
| AQR | PCBP1 | 0 | 0 | 1 | 1 |
| AQR | PTBP1 | 55 | 0.002643937 | 0 | 0.014 |
| AQR | RBM15 | 0 | 0 | 1 | 1 |
| AQR | RBM22 | 0 | 0 | 1 | 1 |
| AQR | SDAD1 | 0 | 0 | 1 | 1 |
| AQR | SRSF1 | 0 | 0 | 1 | 1 |
| AQR | TARDBP | 0 | 0 | 1 | 1 |
| AQR | TIA1 | 5 | 0.000764439 | 0.495 | 0.681 |
| AQR | TRA2A | 0 | 0 | 1 | 1 |
| AQR | U2AF2 | 0 | 0 | 1 | 1 |
| AQR | UCHL5 | 0 | 0 | 1 | 1 |
| AQR | ZC3H11A | 0 | 0 | 1 | 1 |
| AQR | ZNF622 | 0 | 0 | 1 | 1 |
| DROSHA | DROSHA | 14 | 0.002118723 | 1 | 0.8015 |
| DROSHA | EXOSC5 | 19 | 0.007157578 | 0.9355 | 0.0135 |
| DROSHA | GRWD1 | 0 | 0 | 1 | 1 |
| DROSHA | HNRNPC | 0 | 0 | 1 | 1 |
| DROSHA | HNRNPK | 14 | 0.002118723 | 1 | 0.8015 |
| DROSHA | HNRNPM | 19 | 0.003238262 | 1 | 0.999 |
| DROSHA | HNRNPUL1 | 0 | 0 | 1 | 1 |
| DROSHA | HNRNPU | 0 | 0 | 1 | 1 |
| DROSHA | ILF3 | 8 | 0.00131816 | 1 | 0.946 |
| DROSHA | KHSRP | 9 | 0.004106931 | 0.464 | 0.0075 |
| DROSHA | LARP4 | 0 | 0 | 1 | 1 |
| DROSHA | LIN28B | 0 | 0 | 1 | 1 |
| DROSHA | MATR3 | 0 | 0 | 1 | 1 |
| DROSHA | MTPAP | 5 | 0.001119539 | 1 | 0.913 |
| DROSHA | NIPBL | 0 | 0 | 1 | 1 |
| DROSHA | NOLC1 | 0 | 0 | 1 | 1 |

| | | | | | |
|---|---|---|---|---|---|
| DROSHA | PCBP1 | 8 | 0.00131816 | 1 | 0.966 |
| DROSHA | PTBP1 | 0 | 0 | 1 | 1 |
| DROSHA | RBM15 | 12 | 0.001708569 | 1 | 0.783 |
| DROSHA | RBM22 | 0 | 0 | 1 | 1 |
| DROSHA | SDAD1 | 0 | 0 | 1 | 1 |
| DROSHA | SRSF1 | 19 | 0.003238262 | 1 | 1 |
| DROSHA | TARDBP | 0 | 0 | 1 | 1 |
| DROSHA | TIA1 | 17 | 0.006847663 | 0.882 | 0.0045 |
| DROSHA | TRA2A | 0 | 0 | 1 | 1 |
| DROSHA | U2AF2 | 0 | 0 | 1 | 1 |
| DROSHA | UCHL5 | 5 | 0.001119539 | 1 | 0.9995 |
| DROSHA | ZC3H11A | 0 | 0 | 1 | 1 |
| DROSHA | ZNF622 | 0 | 0 | 1 | 1 |
| EXOSC5 | EXOSC5 | 1070 | 0.040277443 | 0 | 0 |
| EXOSC5 | GRWD1 | 0 | 0 | 1 | 1 |
| EXOSC5 | HNRNPC | 0 | 0 | 1 | 1 |
| EXOSC5 | HNRNPK | 21 | 0.007583147 | 0.9565 | 0.012 |
| EXOSC5 | HNRNPM | 1692 | 0.078369367 | 0 | 0 |
| EXOSC5 | HNRNPUL1 | 140 | 0.008026616 | 0 | 0.072 |
| EXOSC5 | HNRNPU | 0 | 0 | 1 | 1 |
| EXOSC5 | ILF3 | 5 | 0.002170825 | 0.983 | 0.3145 |
| EXOSC5 | KHSRP | 57 | 0.008535305 | 0.9985 | 0 |
| EXOSC5 | LARP4 | 0 | 0 | 1 | 1 |
| EXOSC5 | LIN28B | 0 | 0 | 1 | 1 |
| EXOSC5 | MATR3 | 0 | 0 | 1 | 1 |
| EXOSC5 | MTPAP | 14 | 0.001602724 | 0.996 | 0.6585 |
| EXOSC5 | NIPBL | 183 | 0.007020023 | 0.006 | 0.553 |
| EXOSC5 | NOLC1 | 0 | 0 | 1 | 1 |
| EXOSC5 | PCBP1 | 5 | 0.002170825 | 0.992 | 0.391 |
| EXOSC5 | PTBP1 | 0 | 0 | 1 | 1 |
| EXOSC5 | RBM15 | 530 | 0.02722623 | 0 | 0.0005 |
| EXOSC5 | RBM22 | 562 | 0.025448918 | 0 | 0.0055 |
| EXOSC5 | SDAD1 | 0 | 0 | 1 | 1 |
| EXOSC5 | SRSF1 | 67 | 0.014244498 | 1 | 0.0025 |
| EXOSC5 | TARDBP | 277 | 0.016100902 | 0 | 0.0075 |
| EXOSC5 | TIA1 | 65 | 0.006912219 | 0 | 0 |
| EXOSC5 | TRA2A | 0 | 0 | 1 | 1 |
| EXOSC5 | U2AF2 | 0 | 0 | 1 | 1 |
| EXOSC5 | UCHL5 | 832 | 0.040255401 | 0 | 0.0005 |
| EXOSC5 | ZC3H11A | 0 | 0 | 1 | 1 |
| EXOSC5 | ZNF622 | 0 | 0 | 1 | 1 |
| GRWD1 | GRWD1 | 286 | 0.006328706 | 0 | 0.0135 |
| GRWD1 | HNRNPC | 0 | 0 | 1 | 1 |
| GRWD1 | HNRNPK | 0 | 0 | 1 | 1 |
| GRWD1 | HNRNPM | 0 | 0 | 1 | 1 |
| GRWD1 | HNRNPUL1 | 0 | 0 | 1 | 1 |
| GRWD1 | HNRNPU | 0 | 0 | 1 | 1 |
| GRWD1 | ILF3 | 114 | 0.009821615 | 0 | 0.0545 |

| | | | | | |
|---|---|---|---|---|---|
| GRWD1 | KHSRP | 0 | 0 | 1 | 1 |
| GRWD1 | LARP4 | 0 | 0 | 1 | 1 |
| GRWD1 | LIN28B | 0 | 0 | 1 | 1 |
| GRWD1 | MATR3 | 0 | 0 | 1 | 1 |
| GRWD1 | MTPAP | 0 | 0 | 1 | 1 |
| GRWD1 | NIPBL | 0 | 0 | 1 | 1 |
| GRWD1 | NOLC1 | 0 | 0 | 1 | 1 |
| GRWD1 | PCBP1 | 0 | 0 | 1 | 1 |
| GRWD1 | PTBP1 | 0 | 0 | 1 | 1 |
| GRWD1 | RBM15 | 0 | 0 | 1 | 1 |
| GRWD1 | RBM22 | 0 | 0 | 1 | 1 |
| GRWD1 | SDAD1 | 0 | 0 | 1 | 1 |
| GRWD1 | SRSF1 | 468 | 0.017161933 | 0 | 0.0005 |
| GRWD1 | TARDBP | 0 | 0 | 1 | 1 |
| GRWD1 | TIA1 | 0 | 0 | 1 | 1 |
| GRWD1 | TRA2A | 306 | 0.007955296 | 0 | 0.0045 |
| GRWD1 | U2AF2 | 203 | 0.005463766 | 0 | 0.002 |
| GRWD1 | UCHL5 | 350 | 0.008409476 | 0 | 0.9985 |
| GRWD1 | ZC3H11A | 0 | 0 | 1 | 1 |
| GRWD1 | ZNF622 | 7 | 0.000825768 | 1 | 1 |
| HNRNPC | HNRNPC | 69 | 0.000967756 | 0 | 0.952 |
| HNRNPC | HNRNPK | 4 | 0.000282525 | 1 | 1 |
| HNRNPC | HNRNPM | 0 | 0 | 1 | 1 |
| HNRNPC | HNRNPUL1 | 0 | 0 | 1 | 1 |
| HNRNPC | HNRNPU | 14 | 0.001600169 | 0.0025 | 0.5995 |
| HNRNPC | ILF3 | 0 | 0 | 1 | 1 |
| HNRNPC | KHSRP | 0 | 0 | 1 | 1 |
| HNRNPC | LARP4 | 0 | 0 | 1 | 1 |
| HNRNPC | LIN28B | 0 | 0 | 1 | 1 |
| HNRNPC | MATR3 | 0 | 0 | 1 | 1 |
| HNRNPC | MTPAP | 0 | 0 | 1 | 1 |
| HNRNPC | NIPBL | 0 | 0 | 1 | 1 |
| HNRNPC | NOLC1 | 0 | 0 | 1 | 1 |
| HNRNPC | PCBP1 | 0 | 0 | 1 | 1 |
| HNRNPC | PTBP1 | 0 | 0 | 1 | 1 |
| HNRNPC | RBM15 | 0 | 0 | 1 | 1 |
| HNRNPC | RBM22 | 0 | 0 | 1 | 1 |
| HNRNPC | SDAD1 | 0 | 0 | 1 | 1 |
| HNRNPC | SRSF1 | 0 | 0 | 1 | 1 |
| HNRNPC | TARDBP | 0 | 0 | 1 | 1 |
| HNRNPC | TIA1 | 0 | 0 | 1 | 1 |
| HNRNPC | TRA2A | 0 | 0 | 1 | 1 |
| HNRNPC | U2AF2 | 0 | 0 | 1 | 1 |
| HNRNPC | UCHL5 | 0 | 0 | 1 | 1 |
| HNRNPC | ZC3H11A | 0 | 0 | 1 | 1 |
| HNRNPC | ZNF622 | 0 | 0 | 1 | 1 |
| HNRNPK | HNRNPK | 886 | 0.079997389 | 1 | 0.355 |
| HNRNPK | HNRNPM | 19 | 0.003238262 | 1 | 1 |

| | | | | | |
|---|---|---|---|---|---|
| HNRNPK | HNRNPUL1 | 0 | 0 | 1 | 1 |
| HNRNPK | HNRNPU | 81 | 0.001758059 | 0 | 0.921 |
| HNRNPK | ILF3 | 3 | 0.000547356 | 1 | 0.786 |
| HNRNPK | KHSRP | 10 | 0.004373212 | 0.5195 | 0.0085 |
| HNRNPK | LARP4 | 0 | 0 | 1 | 1 |
| HNRNPK | LIN28B | 67 | 0.004905389 | 0.977 | 0.1495 |
| HNRNPK | MATR3 | 0 | 0 | 1 | 1 |
| HNRNPK | MTPAP | 5 | 0.001119539 | 1 | 0.9745 |
| HNRNPK | NIPBL | 0 | 0 | 1 | 1 |
| HNRNPK | NOLC1 | 0 | 0 | 1 | 1 |
| HNRNPK | PCBP1 | 4 | 0.000665195 | 1 | 0.7645 |
| HNRNPK | PTBP1 | 0 | 0 | 1 | 1 |
| HNRNPK | RBM15 | 12 | 0.001708569 | 1 | 0.783 |
| HNRNPK | RBM22 | 0 | 0 | 1 | 1 |
| HNRNPK | SDAD1 | 0 | 0 | 1 | 1 |
| HNRNPK | SRSF1 | 21 | 0.003441596 | 1 | 1 |
| HNRNPK | TARDBP | 0 | 0 | 1 | 1 |
| HNRNPK | TIA1 | 19 | 0.007273231 | 0.8945 | 0.005 |
| HNRNPK | TRA2A | 0 | 0 | 1 | 1 |
| HNRNPK | U2AF2 | 0 | 0 | 1 | 1 |
| HNRNPK | UCHL5 | 5 | 0.001119539 | 1 | 1 |
| HNRNPK | ZC3H11A | 0 | 0 | 1 | 1 |
| HNRNPK | ZNF622 | 0 | 0 | 1 | 1 |
| HNRNPM | HNRNPM | 1293 | 0.053551933 | 0 | 0 |
| HNRNPM | HNRNPUL1 | 185 | 0.012253967 | 0 | 0.004 |
| HNRNPM | HNRNPU | 0 | 0 | 1 | 1 |
| HNRNPM | ILF3 | 12 | 0.002345515 | 1 | 0.9245 |
| HNRNPM | KHSRP | 10 | 0.004373212 | 0.9955 | 0.061 |
| HNRNPM | LARP4 | 0 | 0 | 1 | 1 |
| HNRNPM | LIN28B | 0 | 0 | 1 | 1 |
| HNRNPM | MATR3 | 0 | 0 | 1 | 1 |
| HNRNPM | MTPAP | 28 | 0.002984651 | 1 | 0.7175 |
| HNRNPM | NIPBL | 271 | 0.006450533 | 0 | 0.6075 |
| HNRNPM | NOLC1 | 0 | 0 | 1 | 1 |
| HNRNPM | PCBP1 | 12 | 0.002345515 | 1 | 0.9615 |
| HNRNPM | PTBP1 | 0 | 0 | 1 | 1 |
| HNRNPM | RBM15 | 462 | 0.030964382 | 0 | 0 |
| HNRNPM | RBM22 | 646 | 0.034674357 | 0 | 0 |
| HNRNPM | SDAD1 | 0 | 0 | 1 | 1 |
| HNRNPM | SRSF1 | 117 | 0.011059194 | 1 | 0.6935 |
| HNRNPM | TARDBP | 234 | 0.009461839 | 0 | 0.061 |
| HNRNPM | TIA1 | 40 | 0.01051073 | 0.835 | 0.0005 |
| HNRNPM | TRA2A | 0 | 0 | 1 | 1 |
| HNRNPM | U2AF2 | 0 | 0 | 1 | 1 |
| HNRNPM | UCHL5 | 1232 | 0.070297052 | 0 | 0 |
| HNRNPM | ZC3H11A | 0 | 0 | 1 | 1 |
| HNRNPM | ZNF622 | 0 | 0 | 1 | 1 |
| HNRNPUL1 | HNRNPUL1 | 32 | 0.000850082 | 0.0175 | 0.597 |

| | | | | | |
|---|---|---|---|---|---|
| HNRNPUL1 | HNRNPU | 0 | 0 | 1 | 1 |
| HNRNPUL1 | ILF3 | 0 | 0 | 1 | 1 |
| HNRNPUL1 | KHSRP | 0 | 0 | 1 | 1 |
| HNRNPUL1 | LARP4 | 0 | 0 | 1 | 1 |
| HNRNPUL1 | LIN28B | 0 | 0 | 1 | 1 |
| HNRNPUL1 | MATR3 | 0 | 0 | 1 | 1 |
| HNRNPUL1 | MTPAP | 0 | 0 | 1 | 1 |
| HNRNPUL1 | NIPBL | 0 | 0 | 1 | 1 |
| HNRNPUL1 | NOLC1 | 0 | 0 | 1 | 1 |
| HNRNPUL1 | PCBP1 | 0 | 0 | 1 | 1 |
| HNRNPUL1 | PTBP1 | 0 | 0 | 1 | 1 |
| HNRNPUL1 | RBM15 | 54 | 0.004071155 | 0 | 0.082 |
| HNRNPUL1 | RBM22 | 67 | 0.007262749 | 0 | 0.0695 |
| HNRNPUL1 | SDAD1 | 0 | 0 | 1 | 1 |
| HNRNPUL1 | SRSF1 | 0 | 0 | 1 | 1 |
| HNRNPUL1 | TARDBP | 0 | 0 | 1 | 1 |
| HNRNPUL1 | TIA1 | 0 | 0 | 1 | 1 |
| HNRNPUL1 | TRA2A | 0 | 0 | 1 | 1 |
| HNRNPUL1 | U2AF2 | 0 | 0 | 1 | 1 |
| HNRNPUL1 | UCHL5 | 150 | 0.014611271 | 0 | 0.033 |
| HNRNPUL1 | ZC3H11A | 0 | 0 | 1 | 1 |
| HNRNPUL1 | ZNF622 | 0 | 0 | 1 | 1 |
| HNRNPU | HNRNPU | 67 | 0.001605321 | 0.0315 | 0.9535 |
| HNRNPU | ILF3 | 0 | 0 | 1 | 1 |
| HNRNPU | KHSRP | 0 | 0 | 1 | 1 |
| HNRNPU | LARP4 | 0 | 0 | 1 | 1 |
| HNRNPU | LIN28B | 0 | 0 | 1 | 1 |
| HNRNPU | MATR3 | 0 | 0 | 1 | 1 |
| HNRNPU | MTPAP | 0 | 0 | 1 | 1 |
| HNRNPU | NIPBL | 0 | 0 | 1 | 1 |
| HNRNPU | NOLC1 | 0 | 0 | 1 | 1 |
| HNRNPU | PCBP1 | 0 | 0 | 1 | 1 |
| HNRNPU | PTBP1 | 0 | 0 | 1 | 1 |
| HNRNPU | RBM15 | 0 | 0 | 1 | 1 |
| HNRNPU | RBM22 | 0 | 0 | 1 | 1 |
| HNRNPU | SDAD1 | 0 | 0 | 1 | 1 |
| HNRNPU | SRSF1 | 0 | 0 | 1 | 1 |
| HNRNPU | TARDBP | 0 | 0 | 1 | 1 |
| HNRNPU | TIA1 | 0 | 0 | 1 | 1 |
| HNRNPU | TRA2A | 0 | 0 | 1 | 1 |
| HNRNPU | U2AF2 | 0 | 0 | 1 | 1 |
| HNRNPU | UCHL5 | 0 | 0 | 1 | 1 |
| HNRNPU | ZC3H11A | 0 | 0 | 1 | 1 |
| HNRNPU | ZNF622 | 0 | 0 | 1 | 1 |
| ILF3 | ILF3 | 315 | 0.008804086 | 0 | 0.03 |
| ILF3 | KHSRP | 1 | 0.000601985 | 0.947 | 0.398 |
| ILF3 | LARP4 | 0 | 0 | 1 | 1 |
| ILF3 | LIN28B | 0 | 0 | 1 | 1 |

| | | | | | |
|---|---|---|---|---|---|
| ILF3 | MATR3 | 1 | 0.001001449 | 0.6515 | 0.2205 |
| ILF3 | MTPAP | 4 | 0.001027355 | 0.9905 | 0.2185 |
| ILF3 | NIPBL | 0 | 0 | 1 | 1 |
| ILF3 | NOLC1 | 0 | 0 | 1 | 1 |
| ILF3 | PCBP1 | 4 | 0.000665195 | 1 | 0.7645 |
| ILF3 | PTBP1 | 39 | 0.012290618 | 0 | 0 |
| ILF3 | RBM15 | 7 | 0.001095991 | 1 | 0.9535 |
| ILF3 | RBM22 | 0 | 0 | 1 | 1 |
| ILF3 | SDAD1 | 0 | 0 | 1 | 1 |
| ILF3 | SRSF1 | 119 | 0.014103812 | 0.0015 | 0.0835 |
| ILF3 | TARDBP | 0 | 0 | 1 | 1 |
| ILF3 | TIA1 | 5 | 0.002170825 | 0.9645 | 0.251 |
| ILF3 | TRA2A | 0 | 0 | 1 | 1 |
| ILF3 | U2AF2 | 56 | 0.010663696 | 0 | 0.001 |
| ILF3 | UCHL5 | 57 | 0.004173751 | 0 | 0.0215 |
| ILF3 | ZC3H11A | 0 | 0 | 1 | 1 |
| ILF3 | ZNF622 | 0 | 0 | 1 | 1 |
| KHSRP | KHSRP | 16 | 0.002782838 | 1 | 0.5495 |
| KHSRP | LARP4 | 0 | 0 | 1 | 1 |
| KHSRP | LIN28B | 0 | 0 | 1 | 1 |
| KHSRP | MATR3 | 0 | 0 | 1 | 1 |
| KHSRP | MTPAP | 0 | 0 | 1 | 1 |
| KHSRP | NIPBL | 0 | 0 | 1 | 1 |
| KHSRP | NOLC1 | 0 | 0 | 1 | 1 |
| KHSRP | PCBP1 | 1 | 0.000601985 | 0.963 | 0.4635 |
| KHSRP | PTBP1 | 0 | 0 | 1 | 1 |
| KHSRP | RBM15 | 6 | 0.003146768 | 0.9965 | 0.11 |
| KHSRP | RBM22 | 0 | 0 | 1 | 1 |
| KHSRP | SDAD1 | 0 | 0 | 1 | 1 |
| KHSRP | SRSF1 | 18 | 0.005579725 | 0.9755 | 0.019 |
| KHSRP | TARDBP | 0 | 0 | 1 | 1 |
| KHSRP | TIA1 | 50 | 0.007338782 | 0.952 | 0 |
| KHSRP | TRA2A | 0 | 0 | 1 | 1 |
| KHSRP | U2AF2 | 0 | 0 | 1 | 1 |
| KHSRP | UCHL5 | 0 | 0 | 1 | 1 |
| KHSRP | ZC3H11A | 0 | 0 | 1 | 1 |
| KHSRP | ZNF622 | 0 | 0 | 1 | 1 |
| LARP4 | LARP4 | 24 | 0.004992547 | 1 | 0.992 |
| LARP4 | LIN28B | 0 | 0 | 1 | 1 |
| LARP4 | MATR3 | 0 | 0 | 1 | 1 |
| LARP4 | MTPAP | 0 | 0 | 1 | 1 |
| LARP4 | NIPBL | 0 | 0 | 1 | 1 |
| LARP4 | NOLC1 | 27 | 0.006713325 | 1 | 1 |
| LARP4 | PCBP1 | 0 | 0 | 1 | 1 |
| LARP4 | PTBP1 | 0 | 0 | 1 | 1 |
| LARP4 | RBM15 | 0 | 0 | 1 | 1 |
| LARP4 | RBM22 | 0 | 0 | 1 | 1 |
| LARP4 | SDAD1 | 0 | 0 | 1 | 1 |

| | | | | | |
|---|---|---|---|---|---|
| LARP4 | SRSF1 | 0 | 0 | 1 | 1 |
| LARP4 | TARDBP | 0 | 0 | 1 | 1 |
| LARP4 | TIA1 | 0 | 0 | 1 | 1 |
| LARP4 | TRA2A | 0 | 0 | 1 | 1 |
| LARP4 | U2AF2 | 0 | 0 | 1 | 1 |
| LARP4 | UCHL5 | 0 | 0 | 1 | 1 |
| LARP4 | ZC3H11A | 24 | 0.006223417 | 1 | 0.9745 |
| LARP4 | ZNF622 | 0 | 0 | 1 | 1 |
| LIN28B | LIN28B | 48 | 0.004300985 | 1 | 0.8045 |
| LIN28B | MATR3 | 0 | 0 | 1 | 1 |
| LIN28B | MTPAP | 0 | 0 | 1 | 1 |
| LIN28B | NIPBL | 0 | 0 | 1 | 1 |
| LIN28B | NOLC1 | 0 | 0 | 1 | 1 |
| LIN28B | PCBP1 | 0 | 0 | 1 | 1 |
| LIN28B | PTBP1 | 0 | 0 | 1 | 1 |
| LIN28B | RBM15 | 0 | 0 | 1 | 1 |
| LIN28B | RBM22 | 0 | 0 | 1 | 1 |
| LIN28B | SDAD1 | 0 | 0 | 1 | 1 |
| LIN28B | SRSF1 | 0 | 0 | 1 | 1 |
| LIN28B | TARDBP | 0 | 0 | 1 | 1 |
| LIN28B | TIA1 | 0 | 0 | 1 | 1 |
| LIN28B | TRA2A | 0 | 0 | 1 | 1 |
| LIN28B | U2AF2 | 0 | 0 | 1 | 1 |
| LIN28B | UCHL5 | 0 | 0 | 1 | 1 |
| LIN28B | ZC3H11A | 0 | 0 | 1 | 1 |
| LIN28B | ZNF622 | 0 | 0 | 1 | 1 |
| MATR3 | MATR3 | 437 | 0.058532389 | 0.784 | 0 |
| MATR3 | MTPAP | 0 | 0 | 1 | 1 |
| MATR3 | NIPBL | 0 | 0 | 1 | 1 |
| MATR3 | NOLC1 | 0 | 0 | 1 | 1 |
| MATR3 | PCBP1 | 0 | 0 | 1 | 1 |
| MATR3 | PTBP1 | 567 | 0.066622938 | 0.1545 | 0 |
| MATR3 | RBM15 | 3 | 0.001523753 | 1 | 0.9625 |
| MATR3 | RBM22 | 0 | 0 | 1 | 1 |
| MATR3 | SDAD1 | 0 | 0 | 1 | 1 |
| MATR3 | SRSF1 | 0 | 0 | 1 | 1 |
| MATR3 | TARDBP | 9 | 0.004825877 | 1 | 0.995 |
| MATR3 | TIA1 | 441 | 0.092564276 | 0 | 0 |
| MATR3 | TRA2A | 0 | 0 | 1 | 1 |
| MATR3 | U2AF2 | 0 | 0 | 1 | 1 |
| MATR3 | UCHL5 | 3 | 0.001523753 | 1 | 1 |
| MATR3 | ZC3H11A | 0 | 0 | 1 | 1 |
| MATR3 | ZNF622 | 0 | 0 | 1 | 1 |
| MTPAP | MTPAP | 10 | 0.000764316 | 0.985 | 0.5115 |
| MTPAP | NIPBL | 0 | 0 | 1 | 1 |
| MTPAP | NOLC1 | 0 | 0 | 1 | 1 |
| MTPAP | PCBP1 | 4 | 0.001027355 | 0.9965 | 0.2775 |
| MTPAP | PTBP1 | 0 | 0 | 1 | 1 |

| | | | | | |
|---|---|---|---|---|---|
| MTPAP | RBM15 | 26 | 0.002817568 | 0.9995 | 0.4265 |
| MTPAP | RBM22 | 0 | 0 | 1 | 1 |
| MTPAP | SDAD1 | 0 | 0 | 1 | 1 |
| MTPAP | SRSF1 | 28 | 0.002984651 | 1 | 0.5155 |
| MTPAP | TARDBP | 0 | 0 | 1 | 1 |
| MTPAP | TIA1 | 14 | 0.001602724 | 0.936 | 0.2735 |
| MTPAP | TRA2A | 0 | 0 | 1 | 1 |
| MTPAP | U2AF2 | 0 | 0 | 1 | 1 |
| MTPAP | UCHL5 | 23 | 0.001865112 | 0.982 | 0.2225 |
| MTPAP | ZC3H11A | 0 | 0 | 1 | 1 |
| MTPAP | ZNF622 | 0 | 0 | 1 | 1 |
| NIPBL | NIPBL | 207 | 0.006306817 | 0 | 0.4275 |
| NIPBL | NOLC1 | 0 | 0 | 1 | 1 |
| NIPBL | PCBP1 | 0 | 0 | 1 | 1 |
| NIPBL | PTBP1 | 0 | 0 | 1 | 1 |
| NIPBL | RBM15 | 420 | 0.013874214 | 0 | 0.6185 |
| NIPBL | RBM22 | 223 | 0.006341294 | 0 | 0.8925 |
| NIPBL | SDAD1 | 0 | 0 | 1 | 1 |
| NIPBL | SRSF1 | 1 | 0.000320862 | 1 | 0.9925 |
| NIPBL | TARDBP | 148 | 0.013334235 | 0 | 0.0235 |
| NIPBL | TIA1 | 0 | 0 | 1 | 1 |
| NIPBL | TRA2A | 0 | 0 | 1 | 1 |
| NIPBL | U2AF2 | 0 | 0 | 1 | 1 |
| NIPBL | UCHL5 | 510 | 0.018323364 | 0 | 0.67 |
| NIPBL | ZC3H11A | 0 | 0 | 1 | 1 |
| NIPBL | ZNF622 | 0 | 0 | 1 | 1 |
| NOLC1 | NOLC1 | 61 | 0.018318656 | 1 | 0.6435 |
| NOLC1 | PCBP1 | 0 | 0 | 1 | 1 |
| NOLC1 | PTBP1 | 0 | 0 | 1 | 1 |
| NOLC1 | RBM15 | 0 | 0 | 1 | 1 |
| NOLC1 | RBM22 | 0 | 0 | 1 | 1 |
| NOLC1 | SDAD1 | 0 | 0 | 1 | 1 |
| NOLC1 | SRSF1 | 0 | 0 | 1 | 1 |
| NOLC1 | TARDBP | 0 | 0 | 1 | 1 |
| NOLC1 | TIA1 | 0 | 0 | 1 | 1 |
| NOLC1 | TRA2A | 0 | 0 | 1 | 1 |
| NOLC1 | U2AF2 | 0 | 0 | 1 | 1 |
| NOLC1 | UCHL5 | 0 | 0 | 1 | 1 |
| NOLC1 | ZC3H11A | 24 | 0.012993585 | 1 | 0.782 |
| NOLC1 | ZNF622 | 0 | 0 | 1 | 1 |
| PCBP1 | PCBP1 | 4 | 0.000665195 | 1 | 0.7645 |
| PCBP1 | PTBP1 | 0 | 0 | 1 | 1 |
| PCBP1 | RBM15 | 7 | 0.001095991 | 1 | 0.968 |
| PCBP1 | RBM22 | 0 | 0 | 1 | 1 |
| PCBP1 | SDAD1 | 0 | 0 | 1 | 1 |
| PCBP1 | SRSF1 | 12 | 0.002345515 | 1 | 0.9875 |
| PCBP1 | TARDBP | 0 | 0 | 1 | 1 |
| PCBP1 | TIA1 | 5 | 0.002170825 | 0.978 | 0.3095 |

| | | | | | |
|---|---|---|---|---|---|
| PCBP1 | TRA2A | 0 | 0 | 1 | 1 |
| PCBP1 | U2AF2 | 0 | 0 | 1 | 1 |
| PCBP1 | UCHL5 | 4 | 0.001027355 | 1 | 0.62 |
| PCBP1 | ZC3H11A | 0 | 0 | 1 | 1 |
| PCBP1 | ZNF622 | 0 | 0 | 1 | 1 |
| PTBP1 | PTBP1 | 729 | 0.070570387 | 0 | 0 |
| PTBP1 | RBM15 | 9 | 0.001982772 | 1 | 0.983 |
| PTBP1 | RBM22 | 0 | 0 | 1 | 1 |
| PTBP1 | SDAD1 | 0 | 0 | 1 | 1 |
| PTBP1 | SRSF1 | 0 | 0 | 1 | 1 |
| PTBP1 | TARDBP | 15 | 0.005390903 | 1 | 1 |
| PTBP1 | TIA1 | 520 | 0.115645893 | 0 | 0 |
| PTBP1 | TRA2A | 0 | 0 | 1 | 1 |
| PTBP1 | U2AF2 | 0 | 0 | 1 | 1 |
| PTBP1 | UCHL5 | 11 | 0.002960666 | 1 | 1 |
| PTBP1 | ZC3H11A | 0 | 0 | 1 | 1 |
| PTBP1 | ZNF622 | 2 | 0.000977894 | 1 | 0.9975 |
| RBM15 | RBM15 | 658 | 0.023318913 | 0 | 0.187 |
| RBM15 | RBM22 | 556 | 0.020778611 | 0 | 0.0235 |
| RBM15 | SDAD1 | 0 | 0 | 1 | 1 |
| RBM15 | SRSF1 | 121 | 0.011310815 | 1 | 0.9995 |
| RBM15 | TARDBP | 511 | 0.034651289 | 0 | 0 |
| RBM15 | TIA1 | 33 | 0.008536112 | 0.71 | 0.002 |
| RBM15 | TRA2A | 4 | 0.001479632 | 1 | 1 |
| RBM15 | U2AF2 | 0 | 0 | 1 | 1 |
| RBM15 | UCHL5 | 726 | 0.027249121 | 0 | 0.0015 |
| RBM15 | ZC3H11A | 0 | 0 | 1 | 1 |
| RBM15 | ZNF622 | 93 | 0.007098748 | 0 | 0 |
| RBM22 | RBM22 | 351 | 0.012772844 | 0 | 0.0005 |
| RBM22 | SDAD1 | 0 | 0 | 1 | 1 |
| RBM22 | SRSF1 | 124 | 0.006212017 | 0.444 | 0.5645 |
| RBM22 | TARDBP | 177 | 0.015619144 | 0 | 0.0035 |
| RBM22 | TIA1 | 0 | 0 | 1 | 1 |
| RBM22 | TRA2A | 0 | 0 | 1 | 1 |
| RBM22 | U2AF2 | 0 | 0 | 1 | 1 |
| RBM22 | UCHL5 | 785 | 0.031124682 | 0 | 0.004 |
| RBM22 | ZC3H11A | 0 | 0 | 1 | 1 |
| RBM22 | ZNF622 | 0 | 0 | 1 | 1 |
| SDAD1 | SDAD1 | 0 | 0 | 1 | 1 |
| SDAD1 | SRSF1 | 0 | 0 | 1 | 1 |
| SDAD1 | TARDBP | 0 | 0 | 1 | 1 |
| SDAD1 | TIA1 | 0 | 0 | 1 | 1 |
| SDAD1 | TRA2A | 0 | 0 | 1 | 1 |
| SDAD1 | U2AF2 | 0 | 0 | 1 | 1 |
| SDAD1 | UCHL5 | 0 | 0 | 1 | 1 |
| SDAD1 | ZC3H11A | 0 | 0 | 1 | 1 |
| SDAD1 | ZNF622 | 0 | 0 | 1 | 1 |
| SRSF1 | SRSF1 | 563 | 0.022758621 | 0 | 0.669 |

| | | | | | |
|---|---|---|---|---|---|
| SRSF1 | TARDBP | 37 | 0.007104577 | 0.004 | 0.384 |
| SRSF1 | TIA1 | 57 | 0.012592767 | 0.8725 | 0 |
| SRSF1 | TRA2A | 412 | 0.011112083 | 0 | 0.0005 |
| SRSF1 | U2AF2 | 290 | 0.009461756 | 0 | 0.012 |
| SRSF1 | UCHL5 | 324 | 0.015578123 | 0 | 0.903 |
| SRSF1 | ZC3H11A | 0 | 0 | 1 | 1 |
| SRSF1 | ZNF622 | 15 | 0.00528543 | 0.998 | 0.5905 |
| TARDBP | TARDBP | 421 | 0.017180643 | 0 | 0 |
| TARDBP | TIA1 | 0 | 0 | 1 | 1 |
| TARDBP | TRA2A | 4 | 0.001479632 | 1 | 1 |
| TARDBP | U2AF2 | 0 | 0 | 1 | 1 |
| TARDBP | UCHL5 | 748 | 0.043966412 | 0 | 0 |
| TARDBP | ZC3H11A | 0 | 0 | 1 | 1 |
| TARDBP | ZNF622 | 118 | 0.008674116 | 0.009 | 0.3995 |
| TIA1 | TIA1 | 367 | 0.046289 | 0 | 0 |
| TIA1 | TRA2A | 0 | 0 | 1 | 1 |
| TIA1 | U2AF2 | 0 | 0 | 1 | 1 |
| TIA1 | UCHL5 | 81 | 0.009201278 | 0.1385 | 0.2065 |
| TIA1 | ZC3H11A | 30 | 0.001469929 | 0.0005 | 0.0165 |
| TIA1 | ZNF622 | 0 | 0 | 1 | 1 |
| TRA2A | TRA2A | 828 | 0.02565219 | 0 | 0 |
| TRA2A | U2AF2 | 0 | 0 | 1 | 1 |
| TRA2A | UCHL5 | 732 | 0.028232791 | 0 | 0.7575 |
| TRA2A | ZC3H11A | 0 | 0 | 1 | 1 |
| TRA2A | ZNF622 | 40 | 0.008657224 | 1 | 0.987 |
| U2AF2 | U2AF2 | 73 | 0.001456064 | 0 | 0.8145 |
| U2AF2 | UCHL5 | 30 | 0.001375225 | 0 | 0.2025 |
| U2AF2 | ZC3H11A | 0 | 0 | 1 | 1 |
| U2AF2 | ZNF622 | 0 | 0 | 1 | 1 |
| UCHL5 | UCHL5 | 2199 | 0.074460889 | 0 | 0 |
| UCHL5 | ZC3H11A | 40 | 0.003419587 | 0 | 0.199 |
| UCHL5 | ZNF622 | 203 | 0.011783498 | 0 | 0.0005 |
| ZC3H11A | ZC3H11A | 142 | 0.016680839 | 1 | 0.0005 |
| ZC3H11A | ZNF622 | 0 | 0 | 1 | 1 |
| ZNF622 | ZNF622 | 100 | 0.003991066 | 0 | 0.165 |

**Supplementary Table S2**

| protein 1 | protein 2 | correlations | Summed MI | P (# of correlations) | P (Summed MI) | community 1 | community 2 | | | community 1 | community 2 | observable linkages | connections excluded | % excluded |
|---|---|---|---|---|---|---|---|---|---|---|---|---|---|---|
| PCBP1 | SRSF1 | 12 | 0.002345515 | 1 | 0.9875 | splicing | NS | | | 5' silencing | compartmentalization | 30 | 11 | 37 |
| GRWD1 | ZNF622 | 7 | 0.000825768 | 1 | 1 | 5' silencing | splicing | | | 5' silencing | splicing | 25 | 7 | 28 |
| ILF3 | RBM15 | 7 | 0.001095991 | 1 | 0.9535 | 5' silencing | splicing | | | 5' silencing | U/C | 7 | 0 | 0 |
| NIPBL | SRSF1 | 1 | 0.000320862 | 1 | 0.9925 | 5' silencing | splicing | | | 5' silencing | NS | 18 | 3 | 17 |
| RBM15 | SRSF1 | 121 | 0.011310815 | 1 | 0.9995 | 5' silencing | splicing | | | compartmentalization | splicing | 22 | 2 | 9 |
| RBM15 | TRA2A | 4 | 0.001479632 | 1 | 1 | 5' silencing | splicing | | | compartmentalization | U/C | 2 | 1 | 50 |
| TARDBP | TRA2A | 4 | 0.001479632 | 1 | 1 | 5' silencing | splicing | | | compartmentalization | NS | 9 | 2 | 22 |
| TRA2A | ZNF622 | 40 | 0.008657224 | 1 | 0.987 | 5' silencing | splicing | | | splicing | U/C | 1 | 0 | 0 |
| HNRNPM | PCBP1 | 12 | 0.002345515 | 1 | 0.9615 | 5' silencing | NS | | | splicing | NS | 5 | 1 | 20 |
| LARP4 | ZC3H11A | 24 | 0.006223417 | 1 | 0.9745 | 5' silencing | NS | | | U/C | NS | 2 | 0 | 0 |
| PCBP1 | RBM15 | 7 | 0.001095991 | 1 | 0.968 | 5' silencing | NS | | | 5' silencing | 5' silencing | 32 | 0 | 0 |
| DROSHA | HNRNPM | 19 | 0.003238262 | 1 | 0.999 | 5' silencing | compartmentalization | | | compartmentalization | compartmentalization | 9 | 0 | 0 |
| DROSHA | UCHL5 | 5 | 0.001119539 | 1 | 0.9995 | 5' silencing | compartmentalization | | | splicing | splicing | 11 | 0 | 0 |
| HNRNPK | HNRNPM | 19 | 0.003238262 | 1 | 1 | 5' silencing | compartmentalization | | | U/C | U/C | 1 | 0 | 0 |
| HNRNPK | UCHL5 | 5 | 0.001119539 | 1 | 1 | 5' silencing | compartmentalization | | | NS | NS | 2 | 1 | 50 |
| MATR3 | RBM15 | 3 | 0.001523753 | 1 | 0.9625 | 5' silencing | compartmentalization | | | | | | | |
| MATR3 | TARDBP | 9 | 0.004825877 | 1 | 0.995 | 5' silencing | compartmentalization | | | | | | | |
| MATR3 | UCHL5 | 3 | 0.001523753 | 1 | 1 | 5' silencing | compartmentalization | | | | | | | |
| PTBP1 | RBM15 | 9 | 0.001982772 | 1 | 0.983 | 5' silencing | compartmentalization | | | | | | | |
| PTBP1 | TARDBP | 15 | 0.005390903 | 1 | 1 | 5' silencing | compartmentalization | | | | | | | |
| PTBP1 | UCHL5 | 11 | 0.002960666 | 1 | 1 | 5' silencing | compartmentalization | | | | | | | |
| PTBP1 | ZNF622 | 2 | 0.000977894 | 1 | 0.9975 | 5' silencing | compartmentalization | | | | | | | |
| LARP4 | NOLC1 | 27 | 0.006713325 | 1 | 1 | NS | NS | | | | | | | |
| HNRNPC | HNRNPK | 4 | 0.000282525 | 1 | 1 | compartmentalization | U/C | | | | | | | |
| DROSHA | SRSF1 | 19 | 0.003238262 | 1 | 1 | compartmentalization | splicing | | | | | | | |
| HNRNPK | SRSF1 | 21 | 0.003441596 | 1 | 1 | compartmentalization | splicing | | | | | | | |
| DROSHA | PCBP1 | 8 | 0.00131816 | 1 | 0.966 | compartmentalization | NS | | | | | | | |
| HNRNPK | MTPAP | 5 | 0.001119539 | 1 | 0.9745 | compartmentalization | NS | | | | | | | |

**Supplementary Table S3**

| Gene-specific Primer | Sequence |
|---|---|
| human U1 RT | CAGGGGAAAGCGCGAA |
| human U1 MaP step 1 F | GACTGGAGTTCAGACGTGTGCTCTTCCGATCTNNNNNATACTTACCTGGCAGGG |
| human U1 MaP step 1 R | CCCTACACGACGCTCTTCCGATCTNNNNNCAGGGGAAAGCGCGAA |
| human RMRP RT | ACAGCCGCGCTGAGA |
| human RMRP MaP step 1 F | GACTGGAGTTCAGACGTGTGCTCTTCCGATCTNNNNNGGTTCGTGCTGAAGGC |
| human RMRP MaP step 1 R | CCCTACACGACGCTCTTCCGATCTNNNNNACAGCCGCGCTGAGA |
| human RNase P RT | AATGGGCGGAGGAGAGTAG |
| human RNase P MaP step 1 F | CCCTACACGACGCTCTTCCGATCTNNNNNATAGGGCGGAGGGAAGC |
| human RNase P MaP step 1 R | GACTGGAGTTCAGACGTGTGCTCTTCCGATCTNNNNNAATGGGCGGAGGAGAGTAG |
| mouse U1 RT | TGTGGCAGGGGAGAGCGCG |
| mouse U1 MaP step 1 F | GACTGGAGTTCAGACGTGTGCTCTTCCGATCTNNNNNATACTTACCTGGCAGGG |
| mouse U1 MaP step 1 R | CCCTACACGACGCTCTTCCGATCTNNNNNCAGGGGAGAGCGCG |
| mouse Rmrp RT | GTAGCCGCGCTGAGA |
| mouse Rmrp MaP step 1 F | GACTGGAGTTCAGACGTGTGCTCTTCCGATCTNNNNNAGCTCGCTCTGAAGGC |
| mouse Rmrp MaP step 1 R | CCCTACACGACGCTCTTCCGATCTNNNNNGTAGCCGCGCTGAGA |
| mouse RNase P RT | AGGGCGGGGGAGAGTAG |
| mouse RNase P MaP step 1 F | GACTGGAGTTCAGACGTGTGCTCTTCCGATCTNNNNNAGTGGGCGGAGGAAGC |
| mouse RNase P MaP step 1 R | CCCTACACGACGCTCTTCCGATCTNNNNNAGGGCGGGGGAGAGTAG |

| Reporter Inserts | Sequence |
|---|---|
| pNL3.2.CMV base plasmid | GGCCTAACTGGCCTCAATATTGGCCATTAGCCATATTATTCATTGGTTATATAGCATAAATCAATATTGGCTATTGGCCATTGCATACGTTGTATCTATATCATAATATGTACATTTATATTGGCTCATGTCCAATATGACCGCCATGTTGGCATTGATTATTGACTAGTTATTAATAGTAATCAATTACGGGGTCATTAGTTCATAGCCCATATATGGAGTTCCGCGTTACATAACTTACGGTAAATGGCCCGCCTGGCTGACCGCCCAACGACCCCCGCCCATTGACGTCAATAATGA<br>CGTATGTTCCCATAGTAACGCCAATAGGGACTTTCCATTGACGTCAATGGGTGGAGTATTTACGGTAAACTGCCCACTTGGCAGTACATCAAGTGTATCATATGCCAAGTCCGCCCCCTATTGACGTCAATGACGGTAAATGGCCCGCCTGGCATTATGCCCAGTACATGACCTTACGGGACTTTCCTACTTGGCAGTACATCTACGTATTAGTCATCGCTATTACCATGGTGATGCGGTTTTGGCAGTACACCAATGGGCGTGGATAGCGGTTTGACTCACGGGGATTTCCAAGTCTCC<br>ACCCCATTGACGTCAATGGGAGTTTGTTTTGGCACCAAAATCAACGGGACTTTCCAAAATGTCGTAATAACCCCGCCCCGTTGACGCAAATGGGCGGTAGGCGTGTACGGTGGGAGGTCTATATAAGCAGAGCTCGTTTAGTGAACCGTCAGATCACTAGAAGCTTTATTGCGGTAGTTTATCACAGTTAAATTGCTAACGCAGTCAGTGGGCCTCGGCGGCCAAGCTTGGCAATCCGGTACTGTTGGTAAAGCCACCATGGTCTTCACACTCGAAGATTTCGTTGGGGACTGGCGACAG<br>ACAGCCGGCTACAACCTGGACCAAGTCCTTGAACAGGGAGGTGTGTCCAGTTTGTTTCAGAATCTCGGGGTGTCCGTAACTCCGATCCAAAGGATTGTCCTGAGCGGTGAAAATGGGCTGAAGATCGACATCCATGTCATCATCCCGTATGAAGGTCTGAGCGGCGACCAAATGGGCCAGATCGAAAAAATTTTTAAGGTGGTGTACCCTGTGGATGATCATCACTTTAAGGTGATCCTGCACTATGGCACACTGGTAATCGACGGGGTTACGCCGAACATGATCGACTATTTCGGACGG<br>CCGTATGAAGGCATCGCCGTGTTCGACGGCAAAAAGATCACTGTAACAGGGACCCTGTGGAACGGCAACAAAATTATCGACGAGCGCCTGATCAACCCCGACGGCTCCCTGCTGTTCCGAGTAACCATCAACGGAGTGACCGGCTGGCGGCTGTGCGAACGCATTCTGGCGAATTCTCACGGCTTTCCGCCTGAGGTTGAAGAGCAAGCCGCCGGTACATTGCCTATGTCCTGCGCACAAGAAAGCGGTATGGACCGGCACCCAGCCGCTTGTGCTTCAGCTCGCATCAACGTCTAAGG<br>CGCGACTCTAGAGTCGGGGCGGCCGGCCGCTTCGAGCAGACATGATAAGATACATTGATGAGTTTGGACAAACCACAACTAGAATGCAGTGAAAAAAATGCTTTATTTGTGAAATTTGTGATGCTATTGCTTTATTTGTAACCATTATAAGCTGCAATAAACAAGTTAACAACAACAATTGCATTCATTTTATGTTTCAGGTTCAGGGGGAGGTGTGGGAGGTTTTTTAAAGCAAGTAAAACCTCTACAAATGTGGTAAAATCGATAAGGATCCGTTTGCGTATTGGGCGCTCTTCCGCT<br>GATCTGCGCAGCACCATGGCCTGAAATAACCTCTGAAAGAGGAACTTGGTTAGCTACCTTCTGAGGCGGAAAGAACCAGCTGTGGAATGTGTGTCAGTTAGGGTGTGGAAAGTCCCCAGGCTCCCCAGCAGGCAGAAGTATGCAAAGCATGCATCTCAATTAGTCAGCAACCAGGTGTGGAAAGTCCCCAGGCTCCCCAGCAGGCAGAAGTATGCAAAGCATGCATCTCAATTAGTCAGCAACCATAGTCCCGCCCCTAACTCCGCCCATCCCGCCCCTAACTCCGCCCAGTTCCGCCC<br>TTCTCCGCCCCATGGCTGACTAATTTTTTTTATTTATGCAGAGGCCGAGGCCGCCTCTGCCTCTGAGCTATTCCAGAAGTAGTGAGGAGGCTTTTTTGGAGGCCTAGGCTTTTGCAAAAAGCTCGATTCTTCTGACACTAGCGCCACCATGAAGAAGCCCGAACTCACCGCTACCAGCGTTGAAAAATTTCTCATCGAGAAGTTCGACAGTGTGAGCGACCTGATGCAGTTGTCGGAGGGCGAAGAGAGCCGAGCCTTCAGCTTCGATGTCGGCGGACGCGGCTATGTACTGCGGGTGAA<br>TAGCTGCGCTGATGGCTTCTACAAAGACCGCTACGTGTACCGCCACTTCGCCAGCGCTGCACTACCCATCCCCGAAGTGTTGGACATCGGCGAGTTCAGCGAGAGCCTGACATACTGCATCAGTAGACGCGCCCAAGGCGTTACTCTCCAAGACCTCCCCGAAACAGAGCTGCCTGCTGTGTTACAGCCTGTCGCCGAAGCTATGGATGCTATTGCCGCCGCCGACCTCAGTCAAACCAGCGGCTTCGGCCCATTCGGGCCCCAAGGCATCGGCCAGTACACAACCTGGCGGGATTTCA<br>TTGCGCCATTGCTGATCCCCATGTCTACCACTGGCAGACCGTGATGGACGACACCGTGTCCGCCAGCGTAGCTCAAGCCCTGGACGAACTGATGCTGTGGGCCGAAGACTGTCCCGAGGTGCGCCACCTCGTCCATGCCGACTTCGGCAGCAACAACGTCCTGACCGACAACGGCCGCATCACCGCCGTAATCGACTGGTCCGAAGCTATGTTCGGGGACAGTCAGTACGAGGTGGCCAACATCTTCTTCTGGCGGCCCTGGCTGGCTTGCATGGAGCAGCAGACTCGCTACTTCGAGCC<br>CCGGCATCCCGAGCTGGCCGGCAGCCCTCGTCTGCGAGCCTACATGCTGCGCATCGGCCTGGATCAGCTCTACCAGAGCCTCGTGGACGGCAACTTCGACGATGCTGCCTGGGCTCAAGGCCGCTGCGATGCCATCGTCCGCAGCGGGGCCGGCACCGTCGGTCGCACACAAATCGCTCGCCGGAGCGCAGCCGTATGGACCGACGGCTGCGTCGAGGTGCTGGCCGACAGCGGCAACCGCCGGCCCAGTACACGACCGCGCGCTAAGGAGGTAGGTCGAGTTTAAACTCTAGAACCGG<br>CATGGCCGCAATAAAATATCTTTATTTTCATTACATCTGTGTGTTGGTTTTTTGTGTGTTCGAACTAGATGCTGTCGACCGATGCCCTTGAGAGCCTTCAACCCAGTCAGCTCCTTCCGGTGGGCGCGGGGCATGACTATCGTCGCCGCACTTATGACTGTCTTCTTTATCATGCAACTCGTAGGACAGGTGCCGGCAGCGCTCTTCCGCTTCCTCGCTCACTGACTCGCTGCGCTCGGTCGTTCGGCTGCGGCGAGCGGTATCAGCTCACTCAAAGGCGGTAATACGGTTATCCACAGA<br>ATCAGGGGATAACGCAGGAAAGAACATGTGAGCAAAAGGCCAGCAAAAGGCCAGGAACCGTAAAAAGGCCGCGTTGCTGGCGTTTTTCCATAGGCTCCGCCCCCCTGACGAGCATCACAAAAATCGACGCTCAAGTCAGAGGTGGCGAAACCCGACAGGACTATAAAGATACCAGGCGTTTCCCCCTGGAAGCTCCCTCGTGCGCTCTCCTGTTCCGACCCTGCCGCTTACCGGATACCTGTCCGCCTTTCTCCCTTCGGGAAGCGTGGCGCTTTCTCATAGCTCACGCTGTAGGTATC<br>CAGTTCGGTGTAGGTCGTTCGCTCCAAGCTGGGCTGTGTGCACGAACCCCCCGTTCAGCCCGACCGCTGCGCCTTATCCGGTAACTATCGTCTTGAGTCCAACCCGGTAAGACACGACTTATCGCCACTGGCAGCAGCCACTGGTAACAGGATTAGCAGAGCGAGGTATGTAGGCGGTGCTACAGAGTTCTTGAAGTGGTGGCCTAACTACGGCTACACTAGAAGAACAGTATTTGGTATCTGCGCTCTGCTGAAGCCAGTTACCTTCGGAAAAAGAGTTGGTAGCTCTTGATCCGGCAA<br>ACAAACCACCGCTGGTAGCGGTGGTTTTTTTGTTTGCAAGCAGCAGATTACGCGCAGAAAAAAAGGATCTCAAGAAGATCCTTTGATCTTTTCTACGGGGTCTGACGCTCAGTGGAACGAAAACTCACGTTAAGGGATTTTGGTCATGAGATTATCAAAAAGGATCTTCACCTAGATCCTTTTAAATTAAAAATGAAGTTTTAAATCAATCTAAAGTATATATGAGTAAACTTGGTCTGACAGCGGCCGCAAATGCTAAACCACTGCAGTGGTTACCAGTGCTTGATCAGTGAGGCACCG<br>ATCTCAGCGATCTGCCTATTTCGTTCGTCCATAGTGGCCTGACTCCCCGTCGTGTAGATCACTACGATTCGTGAGGGCTTACCATCAGGCCCCAGCGCAGCAATGATGCCGCGAGAGCCGCGTTCACCGGCCCCCGATTTGTCAGCAATGAACCAGCCAGCAGGGAGGGCCGAGCGAAGAAGTGGTCCTGCTACTTTGTCCGCCTCCATCCAGTCTATGAGCTGCTGTCGTGATGCTAGAGTAAGAAGTTCGCCAGTGAGTAGTTTCCGAAGAGTTGTGGCCATTGCTACTGGCATCGTG<br>GTATCACGCTCGTCGTTCGGTATGGCTTCGTTCAACTCTGGTTCCCAGCGGTCAAGCCGGGTCACATGATCACCCATATTATGAAGAAATGCAGTCAGCTCCTTAGGGCCTCCGATCGTTGTCAGAAGTAAGTTGGCCGCGGTGTTGTCGCTCATGGTAATGGCAGCACTACACAATTCTCTTACCGTCATGCCATCCGTAAGATGCTTTTCCGTGACCGGCGAGTACTCAACCAAGTCGTTTTGTGAGTAGTGTATACGGCGACCAAGCTGCTCTTGCCCGGCGTCTATACGGGACAAC<br>ACCGCGCCACATAGCAGTACTTTGAAAGTGCTCATCATCGGGAATCGTTCTTCGGGGCGGAAAGACTCAAGGATCTTGCCGCTATTGAGATCCAGTTCGATATAGCCCACTCTTGCACCCAGTTGATCTTCAGCATCTTTTACTTTCACCAGCGTTTCGGGGTGTGCAAAAACAGGCAAGCAAAATGCCGCAAAGAAGGGAATGAGTGCGACACGAAAATGTTGGATGCTCATACTCGTCCTTTTTCAATATTATTGAAGCATTTATCAGGGTTACTAGTACGTCTCTCAAGGATAAGTA<br>AGTAATATTAAGGTACGGGAGGTATTGGACAGGCCGCAATAAAATATCTTTATTTTCATTACATCTGTGTGTTGGTTTTTTGTGTGAATCGATAGTACTAACATACGCTCTCCATCAAAACAAAACGAAACAAAACAAACTAGCAAAATAGGCTGTCCCCAGTGCAAGTGCAGGTGCCAGAACATTTCTCT |
| 3' Multiple Cloning Site | CCGCGACTCTAGAGTCG**CTCGAGGGCTGGTACCGTTGCTAGCACGAGCGGCCGCTTACCGGCCGGC**CGCTTCGAGCAGACATG |
| HBA1 intron | GTATGAA**GGTGAGCGGCGGGCCGGGAGCGATCTGGGTCGAGGGGCGAGATGGCGCCTTCCTCTCAGGGCAGAGGATCACGCGGGTTGCGGGAGGTGTAGCGCAGGCGGCGGCTGCGGGCCTGGGCCGCACTGACCCTCTTCTCTGCACA**GGTCTGAGCGG |
| XIST E | AGTCGCTCGAGAAGATCAACATGCCTGGCACTCTAGCACTTGAGGATAGCTGAATGAATGTGTATTTCTTTG**TCTCTTTCTTTCTTGTCTTTGCTCTTTGTTCTCTATCTAAAGTGTGTCTTACCCATTTCCATGTTTCTCTTGCTAATTTCTTTCGTGTGTGCCTTTGCCTCATTTTCTCTTTTTGTTCACAAGAGTGGTCTGTGTCTTGTCTTAGACATATCTCTCATTTTTCATTTTGTTGCTATTTCTCTTTGCTCTCCTAGATGTGGCTCTTCTTTCACGCTTTATTTCATGTC<br>CCTTTTTGGGTCACATGCTGTGTGCTTTTTGTCCTTTTCTTGTTCTGTCTACCTCTCCTTTCTCTGCCTACCTCTCTTTTCTCTTTGTGAACTGTGATTATTTGTTACCCCTTCCCCTTCTCGTTCGTTTTAAATTTCACCTTTTTTCTGAGTCTGGCCTCCTTTCTGCTGTTTCTACTTTTTATCTCACATTTCTCATTTCTGCATTTCCTTTCTGCCTCTCTTGGGCTATTCTCTCTCTCCTCCCCTGCGTGCCTCAGCATCTCTTGCTGTTTGTGATTTTCTATTTCAGTATTAAT<br>TCTGTTGGCTTGTATTTGTTCTCTGCTTCTTCCCTTTCTACTCACCTTTGAGTATTTCAGCCTCTTCATGAATCTATCTCCCTCTCTTTGATTTCATGTAATCTCTCCTTAAATATTTCTTTGCATATGTGGGCAAGTGTACGTGTGTGTGTGTCATGTGTGGCAGAGGGGCTTCCTAACCCCTGCCTGATAGGTGCAGAACGTCGGCTATCAGAGCAAGCATTGTGGAGCGGTTCCTTATGCCAGGCTGCCATGTGAGATGATCCAAGACCAAAACAAGGCCCTAGACTGCAGTAAAA<br>CCAGAACTCAAGTAGGGCAGAAGGTGGAAGGCTCATATGGATAGAAGGCCCAAAGTATAAGACAGATGGTTTGAGACTTGAGACCCGAGGACTAAGATGGAAAGCCCATGTTCCAAGATAGATAGAAGCCTCAGGCCTGAAACCAACAAAAGCCTCAAGAGCCAAGAAAACAGAGGGTGGCCTGAATTGGACCGAAGGCCTGAGTTGGATGGAAGTCTCAAGGCTTGAGTTAGAAGTCTTAAGACCTGGGACAGGACACATGGAAGGCCTAAGAACTGAGACTTGTGACACAAGGCCAAC<br>GACCTAAGATTA**GGTACCGTTGCTAGCACGAGCGGCCGCTTAC |
| XIST 14590-14790 | AGTCGCTCGAG**GATATAGTACTATGCAGACACAAGGAATTGTTTGATAAATGAATAAATTATATGTATATTTGAGGCCAATTTGTGTTTGCTGCTCTGGTAATTTTGAGTAAAAATGCAGTATTCCAGGTATCAGAAACGAAAACACATGGAAACTGCTTTTAAACTTTAAAATATACTGAAAACATAAGGGACTAAGCTTGTTGTGGTCA**GGTACCGTTGCTAGCACGAGCGGCCGCTTAC |
| XIST 18011-18280 | AGTCGCTCGAG**TTAGAAATGTTCTCCTTTTTATCATCAAATCTGCTCAAGAAGGGCTTTTTATAGTAGAATAATATCAGTGGATGAAAACAGCTTAACATTTTACCATGCTTAAGTTTTAAGAATAAAATAAAAATTGGAAATAATTGGCCAAAATTGAAAGGAAAAATTTTTTTAAAATTTCTCTAAATGTAGGCCTGGCTGGGCTTTGACCTTTTCCGTTTTTAAATCACTCACAGAGGGTGGGACAGGAGGAAGAGTGAAGGAAAAGGTCAAACCTGT**GGTACCGTTGCTAGCACGAGCGGCCGCTTAC |
| XIST 18841-18981 | AGTCGCTCGAG**ATGGGGAATGGATTATTGGAAAATGGAAAGAAGAAAGTAACTAAAAGCCTTCCTTTCACAGTTTCTGGCATCACTACCACTACTGATTAAACAAGAATAAGAGAACATTTTATCATCATCTGCTTTATTCACATAAATGAA**GGTACCGTTGCTAGCACGAGCGGCCGCTTAC |
| Control Insert | AGTCGCTCGAC**AACGAGTTGGCGTCACGGAGGCGCAAAAAAGATGATTGTGCCTGCTTCGGCAGTACAACTTCTCGGATAAATGATTATCCGTAACTAGTCCAACTGAATGTTTCCTCGAGTCATAGCTAGCAA**CTAGCACGAGCGGCCGCTTAC |

| qPCR Primer | Sequence |
|---|---|
| pNL3 F | CATCATCCCGTATGAAGGTCTGAGC |
| pNL3 R | TCGATTACCAGTGTGCCATAGTGC |
| 18S F | TTCGAAGACGATCAGATACCGTCG |
| 18S R | CCCGGAACCCAAAGACTTTGG |

Supplemental File 1: RNase P - RMRP structural alignment

```
>RNaseP
--ATAGGGCGGAGGGAAG--CTCATCAG-TGGGGCCACGAGCTGAGTGCGTCCTGTCACTCCACTCCCATGTCCCTTGGG---
AAGGTCTGAGACTAGGGCCAGAGGCGGCCCTAACAGGGCTCTCCCTGAGCTTCGGGGAGGTGAGTTC-----CCAGAGAACGG
GGCTCCGCGCGAGGTC--AGACTGGGCAGGAGATGCCGTGGACCCCGCCCTTCGGGGAGGGGCCCGGCGGATGCCTCCTTTGC
CGGAGCTTGGAACAGACTCACGGCCAGCGAAGTGAGTTC-----AATGGCTGAG---GTGAGGTACCCCGCAGGGGACCTCAT
AACCCAA-----TTCAGACTACTCTCCTCCGCCCATT---

>RMRP
GGTTCGTGCTGA------AGGCCTGTATCCTAGGCTACACACTGAGGAC--TCTGTTCCTCCCCTTTC---CGCCTAGGGGAA
AGTCCCCGGACCTCG----------GG------CAGAGAGTGCCACGTGCATACGCACGTAGA---CATTCCCC---------
-GCTTCCCAC----TCCAAAGTCCGCCAAGAA-----GCGTATCCCGCT-------GAGCGGC-------------GTGGCGC
GGGGGCGT------------CATCC------GTCAGCTCCCTCTAGTTACGCAGGCAGTGCGTGTCCGCGCA---------CC
AACCACACGGGGCTCATTC--------TCAGCGCGGCTGT
```

| Supplemental File 2: eCLIP sites | | |
|---|---|---|
| RBP | XIST start | XIST end |
| AQR | 11943 | 11946 |
| DROSHA | 2076 | 2162 |
| EXOSC5 | 787 | 808 |
| EXOSC5 | 826 | 877 |
| EXOSC5 | 963 | 1042 |
| EXOSC5 | 1077 | 1129 |
| EXOSC5 | 1198 | 1244 |
| EXOSC5 | 1315 | 1393 |
| EXOSC5 | 1435 | 1492 |
| EXOSC5 | 1575 | 1694 |
| EXOSC5 | 1762 | 1822 |
| EXOSC5 | 1850 | 1899 |
| EXOSC5 | 1911 | 1949 |
| EXOSC5 | 2277 | 2350 |
| GRWD1 | 11436 | 11495 |
| GRWD1 | 12962 | 13024 |
| HNRNPC | 450 | 493 |
| HNRNPC | 664 | 694 |
| HNRNPC | 5965 | 6018 |
| HNRNPK | 2056 | 2170 |
| HNRNPK | 2891 | 2974 |
| HNRNPK | 2976 | 3126 |
| HNRNPK | 4256 | 4333 |
| HNRNPK | 4875 | 4947 |
| HNRNPK | 5374 | 5397 |
| HNRNPK | 6076 | 6145 |
| HNRNPK | 6353 | 6414 |
| HNRNPK | 6631 | 6712 |
| HNRNPK | 6942 | 7052 |
| HNRNPK | 7099 | 7134 |
| HNRNPK | 7202 | 7341 |
| HNRNPK | 7480 | 7639 |
| HNRNPK | 7687 | 7733 |
| HNRNPK | 7797 | 7819 |
| HNRNPK | 7822 | 7878 |
| HNRNPK | 8532 | 8582 |
| HNRNPK | 9651 | 9683 |
| HNRNPK | 9738 | 9772 |
| HNRNPK | 9825 | 9889 |
| HNRNPK | 9917 | 10119 |
| HNRNPK | 10127 | 10196 |
| HNRNPM | 914 | 945 |
| HNRNPM | 1005 | 1045 |
| HNRNPM | 1072 | 1121 |
| HNRNPM | 1216 | 1399 |

| | | |
|---|---|---|
| HNRNPM | 1436 | 1493 |
| HNRNPM | 1502 | 1605 |
| HNRNPM | 1754 | 1819 |
| HNRNPM | 1881 | 1924 |
| HNRNPM | 2100 | 2224 |
| HNRNPUL1 | 1445 | 1483 |
| HNRNPU | 4250 | 4287 |
| HNRNPU | 5707 | 5743 |
| HNRNPU | 14669 | 14699 |
| ILF3 | 2061 | 2124 |
| ILF3 | 11634 | 11750 |
| KHSRP | 2326 | 2402 |
| LARP4 | 18829 | 18962 |
| LIN28B | 4904 | 4980 |
| LIN28B | 14521 | 14583 |
| MATR3 | 12036 | 12101 |
| MATR3 | 12134 | 12285 |
| MATR3 | 12288 | 12309 |
| MATR3 | 12331 | 12468 |
| MATR3 | 12481 | 12590 |
| MTPAP | 2198 | 2232 |
| NIPBL | 733 | 780 |
| NIPBL | 815 | 897 |
| NIPBL | 971 | 1029 |
| NOLC1 | 18689 | 18732 |
| NOLC1 | 18754 | 18955 |
| NOLC1 | 18997 | 19070 |
| NOLC1 | 19124 | 19192 |
| PCBP1 | 2069 | 2127 |
| PTBP1 | 11973 | 12035 |
| PTBP1 | 12038 | 12113 |
| PTBP1 | 12136 | 12213 |
| PTBP1 | 12220 | 12320 |
| PTBP1 | 12333 | 12373 |
| PTBP1 | 12380 | 12467 |
| PTBP1 | 12490 | 12536 |
| PTBP1 | 12544 | 12605 |
| PTBP1 | 12640 | 12645 |
| RBM15 | 58 | 81 |
| RBM15 | 115 | 157 |
| RBM15 | 205 | 230 |
| RBM15 | 291 | 402 |
| RBM15 | 405 | 488 |
| RBM15 | 495 | 592 |
| RBM15 | 605 | 635 |
| RBM15 | 651 | 725 |
| RBM15 | 740 | 880 |

| | | |
|---|---|---|
| RBM15 | 895 | 1040 |
| RBM15 | 1215 | 1236 |
| RBM15 | 1333 | 1385 |
| RBM15 | 2094 | 2153 |
| RBM15 | 2186 | 2218 |
| RBM15 | 12718 | 12763 |
| RBM22 | 824 | 902 |
| RBM22 | 997 | 1039 |
| RBM22 | 1185 | 1200 |
| RBM22 | 1312 | 1380 |
| RBM22 | 1638 | 1681 |
| SDAD1 | 494 | 569 |
| SRSF1 | 47 | 106 |
| SRSF1 | 114 | 153 |
| SRSF1 | 201 | 235 |
| SRSF1 | 296 | 398 |
| SRSF1 | 403 | 560 |
| SRSF1 | 604 | 632 |
| SRSF1 | 652 | 781 |
| SRSF1 | 783 | 884 |
| SRSF1 | 2117 | 2256 |
| SRSF1 | 11272 | 11371 |
| SRSF1 | 11436 | 11487 |
| SRSF1 | 12988 | 13053 |
| TARDBP | 1064 | 1144 |
| TARDBP | 12617 | 12754 |
| TIA1 | 2282 | 2334 |
| TIA1 | 12139 | 12218 |
| TIA1 | 12224 | 12291 |
| TIA1 | 12422 | 12469 |
| TIA1 | 13328 | 13395 |
| TRA2A | 12895 | 13088 |
| U2AF2 | 11372 | 11430 |
| UCHL5 | 54 | 94 |
| UCHL5 | 114 | 157 |
| UCHL5 | 197 | 236 |
| UCHL5 | 409 | 441 |
| UCHL5 | 496 | 541 |
| UCHL5 | 652 | 685 |
| UCHL5 | 783 | 807 |
| UCHL5 | 824 | 1049 |
| UCHL5 | 1078 | 1129 |
| UCHL5 | 1146 | 1252 |
| UCHL5 | 1311 | 1316 |
| UCHL5 | 1322 | 1393 |
| UCHL5 | 1585 | 1707 |
| UCHL5 | 1781 | 1794 |

| | | |
|---|---|---|
| UCHL5 | 2179 | 2227 |
| UCHL5 | 11564 | 11573 |
| UCHL5 | 12709 | 12848 |
| UCHL5 | 12966 | 13020 |
| UCHL5 | 13025 | 13085 |
| UCHL5 | 13190 | 13274 |
| ZC3H11A | 13335 | 13356 |
| ZC3H11A | 17763 | 17813 |
| ZC3H11A | 18780 | 18843 |
| ZC3H11A | 18869 | 18931 |
| ZC3H11A | 19148 | 19233 |
| ZNF622 | 12771 | 12853 |
